# Longitudinal single-cell and spatial transcriptomics reveals intratumor heterogeneity, therapeutic response, and comparative value of canine marginal zone lymphoma

**DOI:** 10.64898/2026.09.10.750253

**Authors:** GE Walker, M Macchietto, K Reid, LE Burt, A Winter, S Pracht, M Buettner, V Penza, C Yung, R Dicovitsky, A Kuzmik, DA Vallera, K Demos-Davies, DM Seelig, A Borgatti, M Henson, C Feiock, JF Modiano, S Naik, AL Sarver, AE Treeful

## Abstract

Diffuse B-cell lymphomas are the most prevalent canine hematologic malignancies, and their clinical presentation resembles that of human B-cell non-Hodgkin lymphomas (NHLs). However, a lack of a clear subtyping framework perpetuates imprecise treatment approaches that fail to address mechanisms of therapy resistance. Here, we employed deep phenotyping via serial sampling and longitudinal transcriptomics to evaluate intratumoral composition and therapy response in a 6-year-old neutered male goldendoodle with stage 5A marginal zone lymphoma (MZL). Following sequential treatment consisting of a single IV dose of oncolytic vesicular stomatitis virotherapy (VSV) and standard CHOP chemotherapy 30 days later, we performed scRNAseq on seven serial lymph node biopsies and spatial sequencing on a resection taken 10 days post-VSV treatment. B cells (>90%) comprised three transcriptionally distinct and temporally stable subpopulations, with specific copy number profiles and robust spatial organization despite disrupted lymph node architecture. Analysis across subpopulations revealed downregulation of the quiescence program including *KLF2*, suggesting similarity to *KLF2*-deficient human MZL. While VSV successfully reached target B cells, downstream transcriptional responses were predominantly localized to the T-cell compartment. T cells (≤6%) showed increased proliferation and upregulation of cytotoxicity markers post-VSV administration, enriching for leukocyte-mediated cytotoxicity pathways consistent with an anti-viral immune response. Ultimately, these results provide an *in vivo* proof-of-concept for the treatment paradigm in canine lymphoma, while emphasizing the need for improved subtyping frameworks and multidimensional treatment strategies targeting intratumoral heterogeneity.

**Significance:** Using longitudinal single-cell and spatial transcriptomics, we mapped the molecular architecture and therapeutic response of canine marginal zone lymphoma *in vivo*. We identified conserved *KLF2* downregulation linking canine and human MZL, alongside stable tumor heterogeneity and a distinct immune response to oncolytic virotherapy. These results demonstrate the utility of deep serial phenotyping to track therapy response, while validating naturally occurring canine cancers as comparative models for human therapeutic development.

## Introduction

Diffuse B-cell lymphomas are the most frequent hematological malignant neoplasms in dogs, accounting for 60-70% of all canine lymphomas.^1,2^ Among these, marginal zone lymphomas (MZL) represent a clinically relevant subset whose molecular characteristics remain largely undefined.^3^ In humans, MZL is classified into splenic (SMZL), nodal (NMZL), and extranodal (EMZL/MALT) subtypes based on anatomical site of origin, a stratification that guides prognosis and informs treatment selection.^4,5^ The lack of an analogous subtyping framework in dogs hinders identification of subtype-specific mechanisms driving treatment resistance, thereby creating a critical barrier to achieving durable remissions and impeding the utility of the canine disease as a model for its human counterpart.^6,7^

Oncolytic virotherapy is a promising immunotherapeutic strategy for malignant neoplasms, including canine diffuse B-cell lymphomas. Oncolytic viruses selectively infect tumor cells, inducing immunogenic cell death and release of tumor-associated antigens that stimulate innate and adaptive immune responses. Vesicular stomatitis virus (VSV) is a rapidly replicating RNA virus with demonstrated safety and therapeutic activity in murine tumor models and naturally occurring canine cancer, shown to induce transient remission in dogs with T-cell lymphoma and modest clinical response in dogs with relapsed/refractory B-cell lymphoma by promoting an intratumoral immune response.^8,9^ Oncolytic VSV has been advanced to clinical trials and is additionally being investigated as a neoadjuvant therapy, prior to standard of care tumor resection and chemotherapy protocols.^10^

However, the effect of systemic VSV therapy on the molecular and immune landscape of the canine lymphoma tumor environment has yet to be fully characterized. Single-cell and spatial transcriptomics offer a precise, high-resolution approach to interrogate this, enabling cell type-specific and spatially resolved mapping of tumor composition and therapeutic response. Previous studies have deployed single-cell RNA sequencing (scRNAseq) to investigate intratumoral transcriptional dynamics across tumor and treatment contexts, including oncolytic virotherapy. ^11,12,13,14,15^ Transcriptomic profiling has additionally facilitated the categorization of hematological malignancies, particularly leukemias and lymphoma, into clinically relevant subtypes reflecting their molecular behavior in response to therapy.^16,17,18,19^ Yet the effects of *in vivo* VSV therapy for canine B-cell lymphoma have not yet been evaluated at either the single-cell or spatial level, and detailed molecular stratification of the disease remains uncertain. Here, we employed scRNAseq and spatial transcriptomics to characterize intratumoral cellular heterogeneity of canine MZL and track therapeutic response to oncolytic VSV and subsequent chemotherapy over time. We identified three transcriptionally and spatially distinct B-cell subpopulations exhibiting persistent downregulation of the quiescence program and assessed their copy number variation. We also demonstrate that VSV induces a dynamic anti-viral T-cell response in the tumor within the disrupted architecture of the malignant lymph node. Taken together, these findings provide a molecular framework for understanding clonal evolution in canine MZL and establish the efficacy of VSV to elicit a robust anti-viral immune response leverageable in future therapeutic strategies.

## Materials and Methods

### Study propositus

A six-year-old male neutered Goldendoodle with treatment-naive stage 5A MZL was enrolled in the ORBIT study, designed to investigate combination immunotherapy with an oncolytic VSV and canine PD-1 inhibitor in dogs with advanced lymphoma at the University of Minnesota. The dog was randomly assigned (via www.randomizer.org) to the VSV single-agent arm of the dose-escalation and safety/efficacy phase of the trial. The study was conducted with informed owner consent and under Institutional Animal Care and Use Committee (IACUC)-approved protocols.

### Immunophenotyping and sub-diagnosis

Prior to treatment (day -4), fine needle aspirate (FNA) samples were collected from the left mandibular lymph node for cytologic and flow cytometric evaluation. For cytologic diagnosis, direct smears from the sample were prepared and slides were stained with an aqueous Romanowsky stain (7120 Aerospray hematology slide stainer cytocentrifuge; Wescor). Slides were evaluated by a board-certified veterinary clinical pathologist (DMS) using standard diagnostic techniques. For flow cytometry, samples were collected into a plain red top tube containing flow cytometry buffer with fetal bovine serum (Stain Buffer, BD Pharmingen, Cat. #554656), stained, and analyzed using a previously described approach.^20^ On day 10 post-VSV treatment, the right mandibular lymph node was extirpated, fixed in 10% neutral buffered formalin, and routinely processed and paraffin embedded. From the resultant paraffin block, 5μm histologic sections were stained with hematoxylin and eosin (H&E) or processed for immunohistochemical phenotyping (IHC). For IHC, slides were subjected to antigen retrieval using an EDTA buffer and were incubated with antibodies directed against human CD5, cross-reactive with canine CD5, which is expressed on canine T cells (Neobiotechnologies, clone: C5/473), human CD20, cross-reactive with canine CD20, which is expressed on canine B cells (Thermo Scientific, Cat. RB9013P), human CD79a, cross-reactive with canine CD79a, which is expressed on canine B cells (Biocare Medical, clone: HM47/A9), and human Pax5, cross-reactive with canine Pax5, which is expressed on canine B cells (Neobiotechnologies, clone: PAX5/3977R).

Following antibody incubation, immunodetection was visualized using a dog-appropriate DAKO ENVISION+ kit (Cat. K3468) and the DAB+ chromogen. For each antibody, positive (normal canine lymphoid tissue) and negative (primary antibody replaced with non-immune serum) controls were included. The resultant slides were evaluated for diagnosis and subtyping by a board-certified veterinary pathologist (DMS) according to the canine-adapted version of the World Health Organization’s Classification of Hematopoietic Tumors.^21,22^

### Treatment

On the first day of the trial (day 0), the study dog received a single intravenous dose of 2 × 10^9^ TCID_50_ / kg VSV. Stable disease was maintained for 28 days without further intervention. On day 30 the owners elected to withdraw from the trial to pursue conventional cytotoxic chemotherapy. A standard multi-agent CHOP protocol was initiated as follows: vincristine 0.63-0.7 mg/m^2^ (days 30, 44, 65, 79, 100, 114, 135, and 149), cyclophosphamide 250-270 mg/m^2^ (days 37, 72, 107, and 142), and doxorubicin 25-30 mg/m^2^ (days 51, 86, 121, and 156). A complete response per VCOG-RECIST criteria was achieved at day 100 of the study (70 days post-CHOP initiation), with a five-month duration of response. Following disease progression on day 243, the dog received palliative oral prednisolone at a dose of 1.1 mg/kg q24h, for another four months. The dog was euthanized due to declining quality of life, with 365 days total survival time from study screening.

### Single-cell RNA Sequencing

#### Sample collection

Eight longitudinal lymph node biopsies were collected, including: a pre-treatment (PreTx) Tru-Cut biopsy (day -4); six fine-needle aspirates (FNAs) (days 3, 7, 10, 28, 49, and 70); and a nodal resection (day 10). On treatment days (day 0: VSV; day 49: doxorubicin; day 70: cyclophosphamide), samples were obtained prior to drug administration. Biopsies were collected into RPMI 1640 media (Gibco, Cat. 21870-084) supplemented with 10% fetal bovine serum (FBS, SeraPrime, Cat. F31016-500), Primocin (Invivogen, ant-pm-1), 2 mM L-glutamine (Quality Biological, Cat. QB-118-084-721EA), 2 mM sodium pyruvate (Gibco, Cat. 11360-070), 10 mM 4-(2-hydroxyethyl)-1-piperazineethanesulfonic acid (HEPES, Cat. 25-060-CI), 1% (v/v) MEM Non-Essential Amino Acids Solution (NEAA (100x), Gibco, Cat. 11140-050), and XuM 2-mercaptoethanol (Sigma Aldrich, Cat. M3148), and maintained at 4°C for same-day processing.

#### Preparation of single-cell suspensions

Biopsy samples were mechanically dissociated into single-cell suspensions. Cell concentration and viability were measured by Trypan blue exclusion (Corning, Cat. 25-900-Cl) using a Countess II automated cell counter (Life Technologies). Samples meeting a ≥70% viability threshold were transported on ice to the University of Minnesota Genomics Center for library preparation; the day 70 sample was excluded due to insufficient yield. Upon arrival, cell metrics were reconfirmed using a LUNA-FL™ Automated Cell Counter (Logos Biosystems).

#### Library preparation and sequencing

Single-cell capture and library preparation were performed according to the 10x Genomics Chromium NEXT GEM Single Cell 3’ Gene Expression (v3.1) protocol. cDNA quality and size distribution were assessed via Agilent Bioanalyzer (Agilent, Santa Clara, CA), and quantification was performed using a Qubit Fluorometer (Thermo Fisher Scientific, Waltham, MA). Dual-indexed libraries were sequenced on an Element Aviti platform (Element Biosciences, San Diego, CA). Raw FASTQ files for all 7 samples were aligned to a custom reference build comprising the canFam4 genome (NCBI RefSeq, accession GCF_011100685.1), the canine Y chromosome (NBCI RefSeq, accession NC_051844.1), and the canine mitochondrial genome (Ensembl release 104).^23,24^ Alignment was performed with CellRanger (v8.0.1) to generate expression matrices from gene barcodes.

#### Quality control, filtering, and clustering

Gene expression matrices were converted into 7 individual Seurat objects using the Seurat package (v5.3.0) in R (v4.4.3). Objects were first processed independently to assess sample quality and to remove low-quality or dying cells. Cells were filtered at sample-specific thresholds to remove those with high mitochondrial expression and low feature/UMI counts. PreTx, day 7, day 28, and day 49 showed bimodal count distributions indicative of a population of low-quality contaminant cells, which were subsequently removed. Genes expressed in less than 10 cells were also removed. DoubletFinder (v2.0.4) was used to identify potential doublet captures, and predicted barcodes were filtered out.

Following independent processing, samples were merged into a single Seurat object and normalized with SCTransform. Principal component analysis was performed, and an elbow plot determined an optimal 25 PCs for input into neighbor’s graph generation. Cells were clustered with the Louvain-graph based clustering algorithm at various resolutions. An ideal resolution of 0.05 was determined by plotting the varying cluster cell assignments with the clustree package (v0.5.1). Cell clusters were visualized using uniform manifold approximation and projection (UMAP) plots.

#### Cell annotation and typing

Cell lineages were inferred through a combination of cluster differentially expressed genes (DEGs), cell type marker expression, and automated annotation with SingleR (v2.8.0). The SingleR cell type assignments were predicted using 4 built in reference datasets from celldex (v1.16.0), in lieu of an existing reference canine dataset: BlueEncode, ImmGen, and both main and fine labels of the Human Primary Cell Atlas Data. Final automated cell types were determined as those which were consensus across all four reference annotations. Top cluster DEGs were calculated with Wilcoxon rank-sum tests using the FindAllMarkers() function in Seurat, with a p_adj cutoff of <0.05. Expression of cell type markers were visualized with feature plots and violin plots. Cell type predictions were referenced against cluster DEGs and cell type markers to yield final annotations.

#### T cell subclustering and G2M signature expression

Cells belonging to the initial T-cell cluster were extracted, renormalized, and reclustered using the same methodology as above. An optimal resolution of 0.06 was iteratively determined again by comparing clustering at various resolutions with the clustree package. Clusters were annotated using the same three-dimensional consensus approach as before of cluster DEGs, cell type markers, and automated cell label predictions.

G2M signature expression was calculated on the subset T cell population using the function AddModuleScore() with the literature-curated cell cycle marker gene set loaded with Seurat.^25^ The resulting module score expression was plotted as both a UMAP, and as a boxplot across all T cells from the original clustering between PreTx and day 10 resection. Differences in G2M module scores between timepoints were statistically assessed at the single-cell level using a Wilcoxon rank-sum test.

#### DGE and pathway analysis

Differential gene expression (DGE) testing was performed using the Seurat FindMarkers() function with a non-parametric Wilcoxon rank-sum test. To mitigate inflated statistical results driven by large cell numbers, 500 cells were randomly sampled per timepoint for both B cells and non-malignant lymphocytes, across all pairwise time comparisons. Differential genes for select timepoint pairs (*e.g.*, PreTx and day 3) were initially filtered at a standard log2_FC of 0.6 and a pval_adj cutoff of 0.05, and successively filtered again at increasing stringent cutoffs until < 500 DEGs remained. The filtered DEG list was then input into ClusterProfiler (v4.14.6) for Over-Representation Analysis (ORA), with a GS cutoff of 5 and a pval_adj cutoff of 0.15 for enriched pathways. Differentially expressed genes were visualized with the EnhancedVolcano (v1.24.0) package in R.

#### Detection of viral RNA transcripts

To test for VSV transcripts in the single-cell data, raw FASTQ files for each timepoint were aligned to the VSV genome downloaded from NCBI (GenBank: FJ478454.1) with bowtie2 (v4.8.5). The alignment was repeated with the canine CD19 (XM_038668761.1) and GAPDH (NM_001003142.2) genes to provide a positive control.

#### VSV qRT-PCR

Virus pharmacokinetics and viremia were assessed by qRT-PCR of whole blood (WB) and FNAs from lymph nodes. Approximately 500 μL of WB was collected in RNAprotect® Animal Blood Tubes (Qiagen Cat. 76554) at baseline (pre-treatment), 6 hours (day 0) and 24h (day 1) post-VSV administration, followed by RNA isolated with RNeasy Protect Animal Blood kit (Qiagen Cat. 73224). Serial FNAs were collected into sterile PBS, centrifuged to obtain cell pellets, and the supernatant was discarded.

Pellets were flash-frozen in liquid nitrogen and stored at -80° C. qRT-PCR for the detection of VSV-N was performed with LightCycler ® 480 RNA master Hydrolysis Probes kit (Roche Cat. 04707494001) according to protocol. Forward primer 5’-TGATAGTACCGGAGGATTGACGAC-3’, reverse primer 5’-CCTTGCAGTGACATGACTGCTCTT-3’ and probe 5’-FAM-TCGACCACATCTCTGCCTTGTGGCGGTGCA-BHQ1-3’ were designed to amplify both negative sense genomic VSV RNA and positive sense mRNA VSV-N. qRT-PCR reaction was performed on a QuantStudio7 Pro instrument. Relative quantification of VSV-N copies per 1 ug of isolated RNA was obtained by running VSV-N plasmid standard curve alongside samples.

### Spatial Transcriptomics

#### Sample collection and processing

A 1cm^2^ nodal section was flash-frozen in a pre-chilled isopentane/dry ice bath, embedded in O.C.T. compound (Tissue-Tek®, Cat. 4583), and stored at -80°C. Tissue quality was confirmed with a RIN score of 7.5, and 10-µm sections were mounted on Visium capture slides. Following the 10X Genomics Tissue Optimization protocol, an 18-minute permeabilization time was identified as optimal. H&E staining and imaging were performed on a Huron TissueScope^TM^ LE prior to library preparation. Library quality and quantification were confirmed via Agilent Bioanalyzer and Qubit. Dual-indexed libraries were sequenced on an Element Aviti platform and FASTQs were aligned to the Canfam4_Y reference canine genome with SpaceRanger (v4.0.1).

#### Quality control and preprocessing

SpaceRanger output files were converted to AnnData objects and preprocessed in Python using Scanpy (v1.11.5) and Squidpy (v1.6.5). A total of 333 spots (2.3%) expressing fewer than 250 unique genes, 250 total counts, or exhibiting greater than 10% mitochondrial expression were removed, yielding 4,601 high-quality spots for analysis. Data were normalized, log-transformed, and the top 3,000 highly variable genes were selected for PCA and clustering.

An optimal 25 PCs were used to generate UMAP embeddings. Leiden clustering was performed at various resolutions, and an optimal resolution determined by a combination of ASW scores per clustering and visual inspection. Cluster DEGs were calculated using Wilcoxon rank-sum tests and visualized with dot plots. Marker genes were plotted both as UMAPs and as spatial plots to visually examine expression.

#### Spot-based deconvolution

As the spot-based resolution of the data limited successful manual annotation of cell types, deconvolution was performed to transfer labels from a single-cell reference and achieve more refined annotation. Cell2location (v0.1.5) was employed, using the day 10 single-cell RNAseq dataset as a reference to acquire cell type gene signatures and subsequently map them to the spatial dataset. The model was generated with detection_alpha=20 and N_cells_per_location=8 in concordance with Visium spot-based data, and training was conducted over 15,000 epochs. Resulting predicted cell type proportions per each 50 µm spot were plotted spatially.

#### Spatially variable gene analysis

Spatially variable genes were calculated in SquidPy using Moran’s I global spatial auto-correlation statistics. Genes were scored based on the extent to which their expression shows a pattern that is spatially clustered throughout the tissue, as opposed to randomly dispersed. The top 20 highest scoring genes were plotted spatially to visualize expression.

#### Composition-based clustering

The R package ISCHIA was employed to perform cell-type composition-based clustering of spatial spots. The AnnData object was first converted into a Seurat object for compatibility. A deconvoluted cell type probability matrix was generated from cell2location predictions, normalized across spots, and an optimal k=5 determined using an elbow plot. Clustering was performed using the Composition.cluster() function and resulting cluster assignments were plotted spatially. Box plots displaying cell type abundances for each composition cluster were generated using the Composition_cluster_enrichedCelltypes() function.

### Statistical Analysis

For identifying cluster markers and differentially expressed genes in scRNAseq data, Wilcoxon rank-sum tests were employed with a Benjamini–Hochberg correction for calculation of adjusted p-values. Over representation analysis with ClusterProfiler used a hypergeometric test to identify enriched pathways with a Benjamini-Hochberg correction for calculation of adjusted p-values. For Moran’s I spatial autocorrelation, statistical significance was determined using permutation-based testing. Resulting p-values were corrected for multiple hypothesis testing using the Benjamini–Hochberg false discovery rate (FDR) method.

## Data Availability

The datasets generated and analyzed in this study are publicly available through the NCBI Gene Expression Omnibus (GEO) under accession number GSE343505.

## Code Availability

All underlying code for the analyses in this study can be found on GitHub at https://github.com/walk1195/longitudinal-canine-mzl.

## Results

### Cytology, flow cytometry, and IHC confirm an immunophenotype consistent with MZL

The FNA cytology samples were diagnostic for lymphoma through the identification of a dominant population of intermediate to large lymphocytes measuring 1.25 to 1.5x the diameter of a neutrophil, which contained scant to small amounts of basophilic cytoplasm and an often-paracentral nucleus with finely stippled chromatin and a single nucleolus (**Fig. 1A**). The flow cytometry study identified a population of CD21+ B cells that represented 94.9% of all cellular events and were intermediate-sized with a median forward scatter measuring 1.5x the median forward scatter of this dog’s CD5+ T cells (**Fig. 1B-C**). Review of the H&E-stained slide revealed diffuse effacement of the lymph node architecture by a near-uniform population of intermediate-sized lymphocytes containing a small amount of pink to light blue cytoplasm and a round to slightly ovoid, centrally-located nucleus measuring 1.25 to 1.5x the diameter of a red blood cell (**Fig. 1D**). Nuclear chromatin was finely granular to smooth and 5 mitotic figures were seen in 10, 400x fields. Approximately 90-95% of these cells demonstrated immunopositivity for CD20, Pax5, and CD79a whereas only 10% demonstrated immunopositivity for CD5 (**Fig. 1E-F**). In aggregate, these findings were diagnostic for non-large cell, diffuse B-cell lymphoma, and based upon the neoplastic cellular features of a round to oval nuclear shape, dispersed chromatin, low mitotic index, and centrally-located nucleolus, most consistent with canine NMZL.

**Figure 1.**
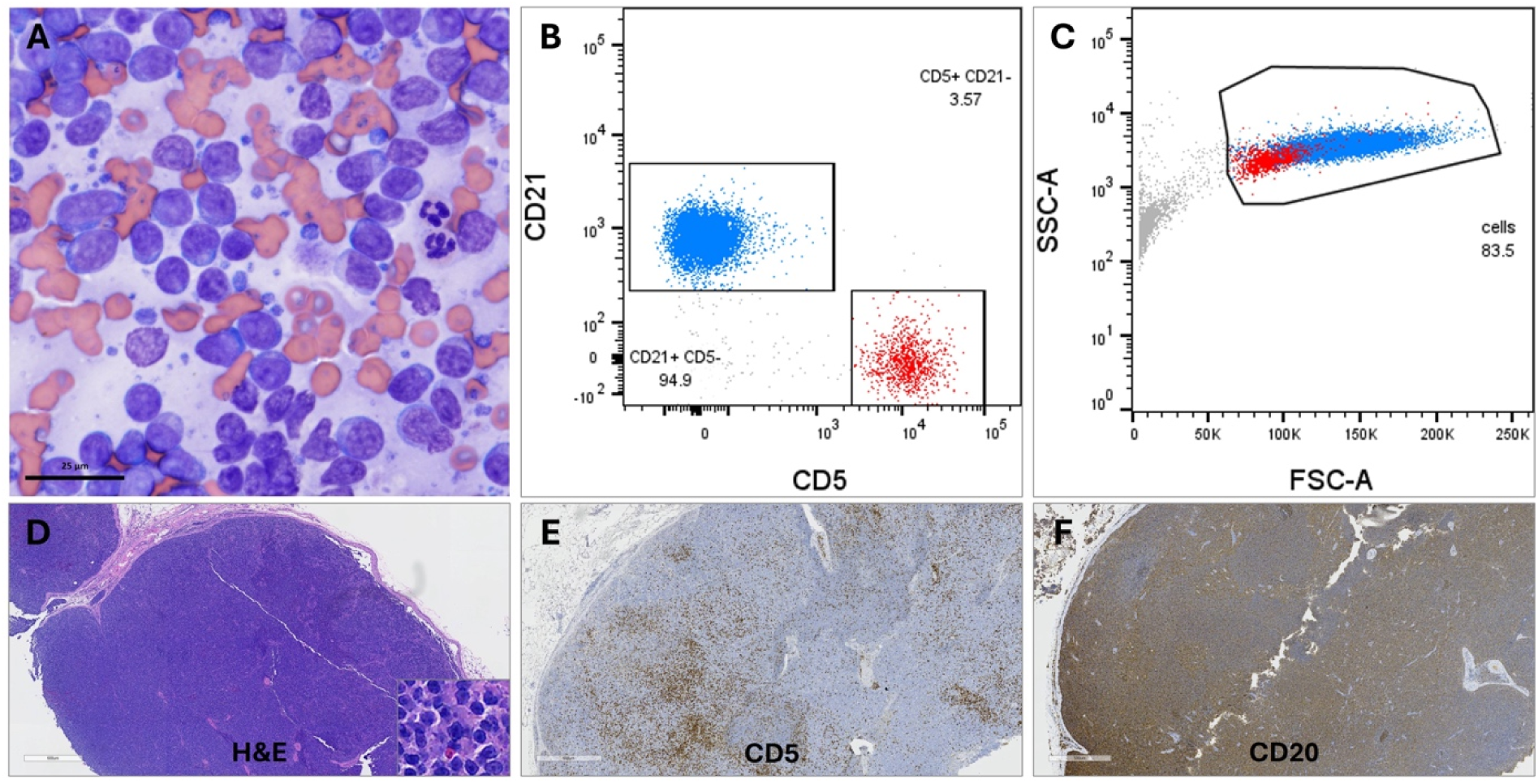
Cytology, flow cytometry, and IHC confirm an immunophenotype consistent with MZL. **A)** Fine needle aspirate cytology of an enlarged lymph node demonstrates a near homogenous population of intermediate-sized lymphocytes. **B-C)** Flow cytometric analysis of the same lymph node reveals 94.9% CD21+ B cells (labelled blue) that demonstrate larger median forward scatter as compared to this dog’s CD5+ T cells (labelled red). **D-F)** Histopathologic review of an extirpated lymph node demonstrates an effacing, intermediate-sized (inset) lymphoid population that is CD5 (brown) immunonegative and CD20 (brown) immunopositive.

### VSV treatment does not induce alterations in three transcriptionally distinct B-cell lymphoma subpopulations

We hypothesized that selective pressure introduced by VSV infection would drive clonal evolution of the tumor cells. To test this, we characterized the clonal architecture of the tumor in the propositus to gain insight to its cellular composition and transcriptional dynamics. Following independent filtering of each serial sample passing QC (n=7), a total of 176,143 high-quality cells were retained for downstream analysis (**Fig. 2A**).

**Figure 2.**
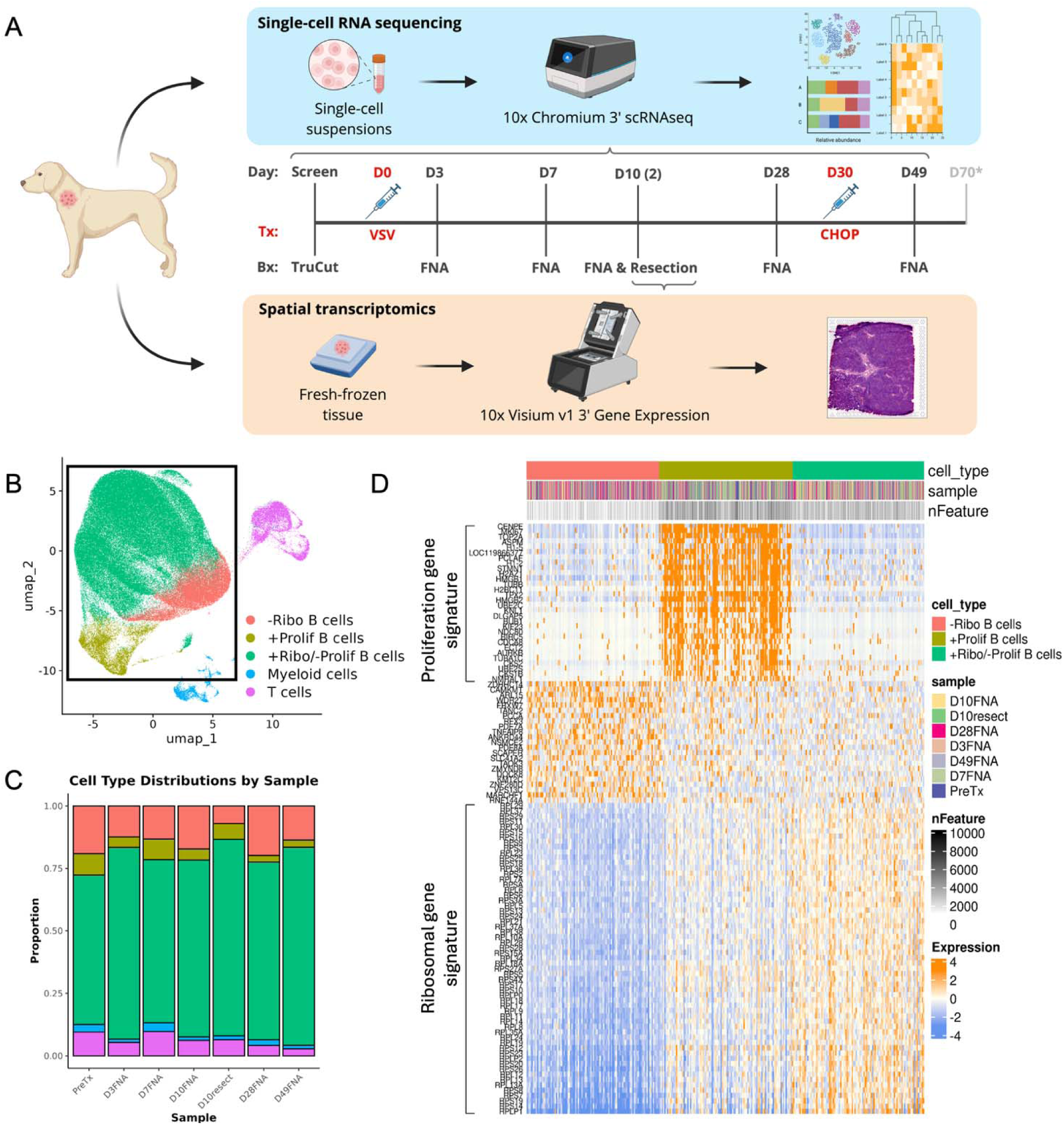
VSV treatment does not induce alterations in three transcriptionally distinct B-cell lymphoma subpopulations. **A)** Overview of sampling and sequencing time course following VSV treatment administration. scRNA-seq was performed on longitudinal biopsies and spatial transcriptomics on the day 10 resection. **B)** UMAP of cell populations present in the tumor. Cells from all time points were merged and plotted together; 3 unique B-cell subpopulations are highlighted by a black box. **C)** Bar plot of cell type distributions across timepoint samples. **D)** Heatmap of unique gene signature expression across the 3 B-cell subpopulations (500 randomized cells each) shown in B.

Sample quality was comparable across timepoints, with the day 10 resection yielding the most uniform sequencing depth and lowest proportion of low-quality cells **(Supplemental Fig. S1A-C)**. Clustering of cells across all timepoints in aggregate revealed a large B-cell population present with minimal batch effects, expressing canonical B cell markers (*e.g. CD19, CD20, CD79A*) (**Fig. 2B, Supplemental Fig. S1D**). A separate population of enriched cells expressed T-cell markers (*e.g., CD3E*, *CD3G, CD3D*), and another showed expression indicative of myeloid cells (*e.g., CD163, CD68*) **(Supplemental Fig. S2A)**. Cell type annotations were validated by automated prediction methods, reinforcing confidence in the assigned labels **(Supplemental Fig. S2B,C**).

Within the B-cell compartment, analysis revealed transcriptionally unique expression profiles defining three distinct subpopulations (**Fig. 2C**). Approximately 5.6% of all B cells expressed a signature that was enriched for genes associated with cell cycle progression and the DNA damage response, such as *TOP2A, MKI-67, CENPF, STMN1, HMGB1, HMGB2, TPX2, BUB1,* and *H2AZ1*. A second subpopulation (∼16.6%) showed downregulation of ribosomal genes (*RPS19, RPS26, RPLP1, RPS14, RPL9*), while exhibiting low mitochondrial expression. The third and largest B-cell subpopulation (∼77.8%) was uniquely characterized by a hybrid state, lacking differential expression of proliferation and DNA damage response genes but possessing a detectable ribosomal program. Only slight variability in the proportions of each cell population was appreciated across the longitudinal timepoints, indicating the clonal architecture was lineage specific and largely unaffected by VSV-driven selection (**Fig. 2C**). Further validation with GCESS gene-scoring analysis confirmed this temporal stability of cell populations across the sampling time course (**Supplemental Fig. S2D**).^26^

### B-cell subpopulations possess detectable copy number alterations

To further interrogate the clonal relationship of the three B-cell subpopulations within the tumor, we inferred copy number gains and losses using T cells and myeloid cells as references. The ribosomal-depleted B cells exhibited the most consistently aberrant copy number variation (CNV) profile, including a predicted loss at chromosome 20 (CFA20), the genomic location of *KLF2* (**Fig. 3A**). This pattern was uniformly predicted across all cells in the subpopulation and consistently present across all timepoints, suggesting possible clonal expansion of these B-cells (**Supplemental Fig. S3**). Notably, a subset of +Ribo/–Prolif B cells shared several of the predicted CNV gains (e.g., CFA11, CFA12, CFA14, CFA22, CFA32, CFA36) (**Fig. 3A**). A separate, smaller subset of the +Ribo/-Prolif B cells displayed a predicted gain on CFA13, the site of the *MYC* and *KIT* oncogenes and a hallmark of ∼40% of canine non-Hodgkin lymphomas (**Fig. 3A**).^27,28^ Together these CNV patterns highlight genomic heterogeneity within the tumor, suggesting partially overlapping clonal relationships, likely from a common origin that require further molecular subtyping to fully contextualize.

**Figure 3.**
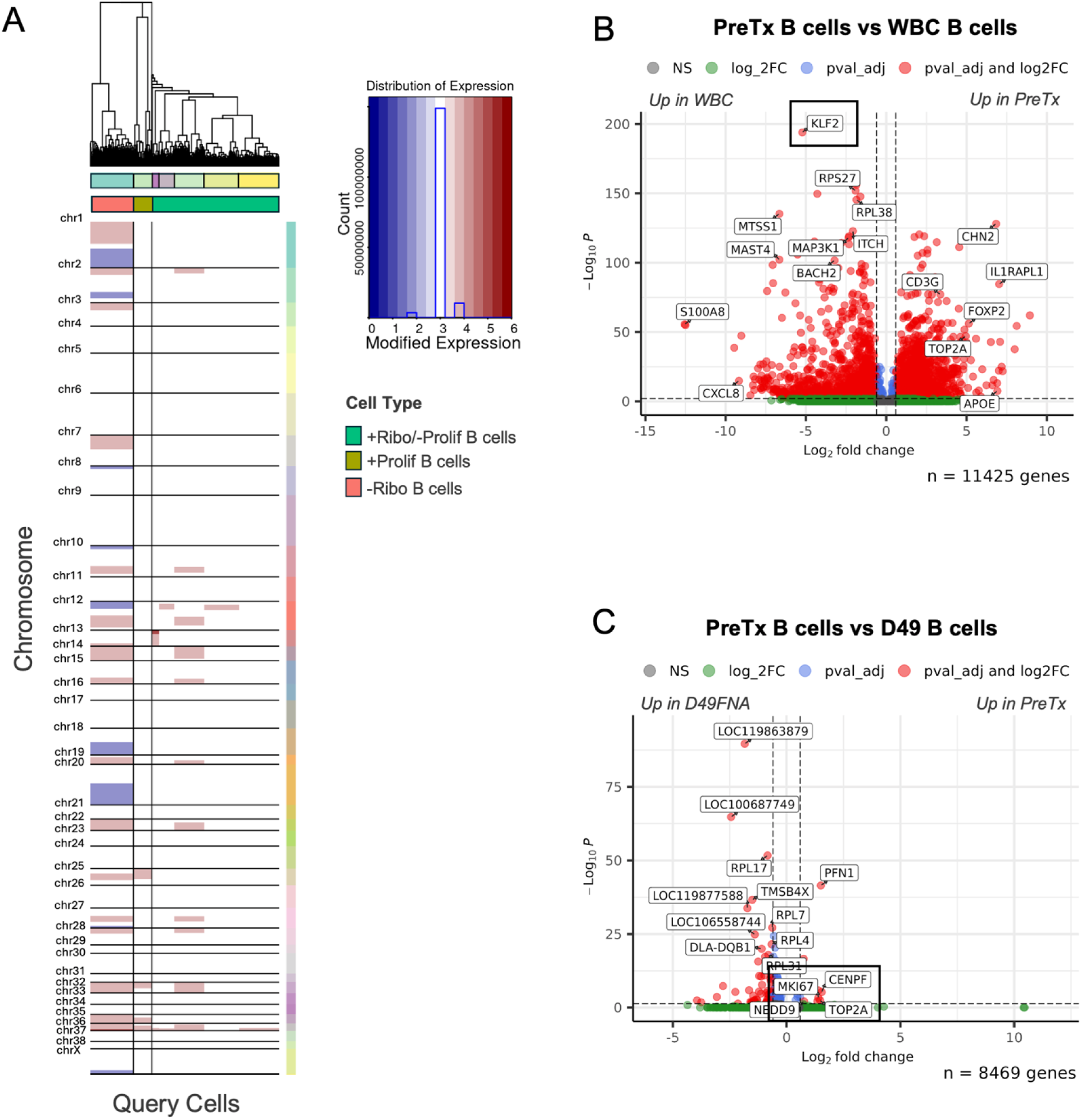
MZL B cells exhibit copy number alterations, persistent downregulation of the quiescence program, and elimination of the proliferative pool after chemotherapy. **A)** Denoised heatmap predicted CNVs across the three B-cell subpopulations, using T cells and myeloid cells as the reference cell type. **B)** Volcano plot of differentially expressed genes between PreTx sample B cells and B cells from PBMCs of a healthy dog. *KLF2* is indicated by a black box. **C)** Volcano plot showing downregulation of proliferation marker genes *TOP2A*, *MKI67*, and *CENPF* post-chemotherapy treatment.

### MZL B cells exhibit persistent downregulation of the quiescence program

We next investigated how the transcriptional landscape of the MZL B cell population differed from normal peripheral B cells. DGE analysis revealed dysregulation of numerous genes in the PreTx B cells, including downregulation of *BACH2*, *MAST4*, *MAP3K1*, and *RIPOR2*, and upregulation of *FOXP2*, *TOP2A*, *CD3G*, and *IL1RAPL1* (**Fig. 3B**). Notably, *KLF2* was the most highly downregulated gene in the PreTx B cells. *KLF2*, a tumor suppressor and regulator of inflammatory gene activation in humans, has been implicated as the most frequently mutated gene in human SMZL, providing an intriguing parallel between human and canine MZLs.^29^ The downregulation of *BACH2* and *RIPOR2* further suggests that compared to normal peripheral B cells, the MZL B cells exhibit transcriptional downregulation of their quiescence program.

### Early viral replication in the tumor microenvironment precedes chemotherapy-induced depletion of the proliferating cell pool

Next, we examined how VSV and CHOP treatments affected the tumor’s B cell population at the gene expression level, hypothesizing that VSV infection of tumor cells would generate detectable viral RNA transcripts and/or B-cell depletion, while CHOP would suppress tumor-cell growth, eliminating the proliferation signature. While VSV transcripts were not captured via scRNAseq, likely due to transcript levels below the detectable threshold, targeted qRT-PCR demonstrated robust VSV-N copy numbers at day 3, confirming successful delivery and active viral replication within the tumor microenvironment (**Supplemental Fig. S4**). This early replication occurred, leaving the broader host B-cell transcriptome largely unaltered throughout the pre-chemotherapy window. (**Supplemental Fig. S5A**). Predictably, proliferation genes *MKI67*, *CENPF*, and *TOP2A* showed downregulation in the day 49 (post-CHOP) sample when compared to the PreTx sample (**Fig. 3C**). The day 49 B cells also showed downregulation of canonical markers *CD79A*, *CD79B*, and *JCHAIN*, of ribosomal genes, and of ATP synthase subunits compared to B cells from the day 3 and day 28 samples (**Supplemental Fig. S5B-E**). The ribosomal genes showed enrichment for pathways implicated in ribosome assembly and translation (**Supplemental Fig. S5B-E**). Together, these data indicate that CHOP chemotherapy successfully eliminated metabolically active, proliferating cells in the tumor.

### VSV induces a functionally intact anti-viral T-cell response despite a disrupted lymph node architecture

To test the hypothesis that VSV exposure primes and activates T cells, we reclustered and reannotated the T cell population (**Fig. 4A**). Reprocessing identified 6 subpopulations, including CD4+ cells (*CD4, CD3G, CD3D*), CD8+ cells (*CD8A, CD8B*), and a putative NK cell population expressing *NKG7, GMZA, and CD160* (**Supplemental Fig. S6A,B**). A subset showed expression of interleukin-23 receptor (*IL-23R*) and *IL1R1*, subsequently referred to as IL23R+ T cells (**Supplemental Fig. S6C**). Another subpopulation showing strong expression of cell cycle markers *MKI67*, *CENPF*, and *TOP2A* consistent with T-cell proliferation was labelled as proliferating T cells (**Fig. 4A, Supplemental Fig. S6A,B**). Cells denoted as naïve T cells included conventional CD4+ cells, CD8*+* cells, and CD4-CD8-cells, showing upregulation of *FOXP1*, *RUNX1*, and *KLF12* indicative of transcriptional restraint and quiescence. A small population of cells was annotated as naïve B cells. Close analysis of their transcriptional profile indicated that these cells most likely represented a population of residual, non-malignant B cells in the sampled lymph node, whose gene expression patterns more closely resembled non-malignant T cells than malignant B-cells.

**Figure 4.**
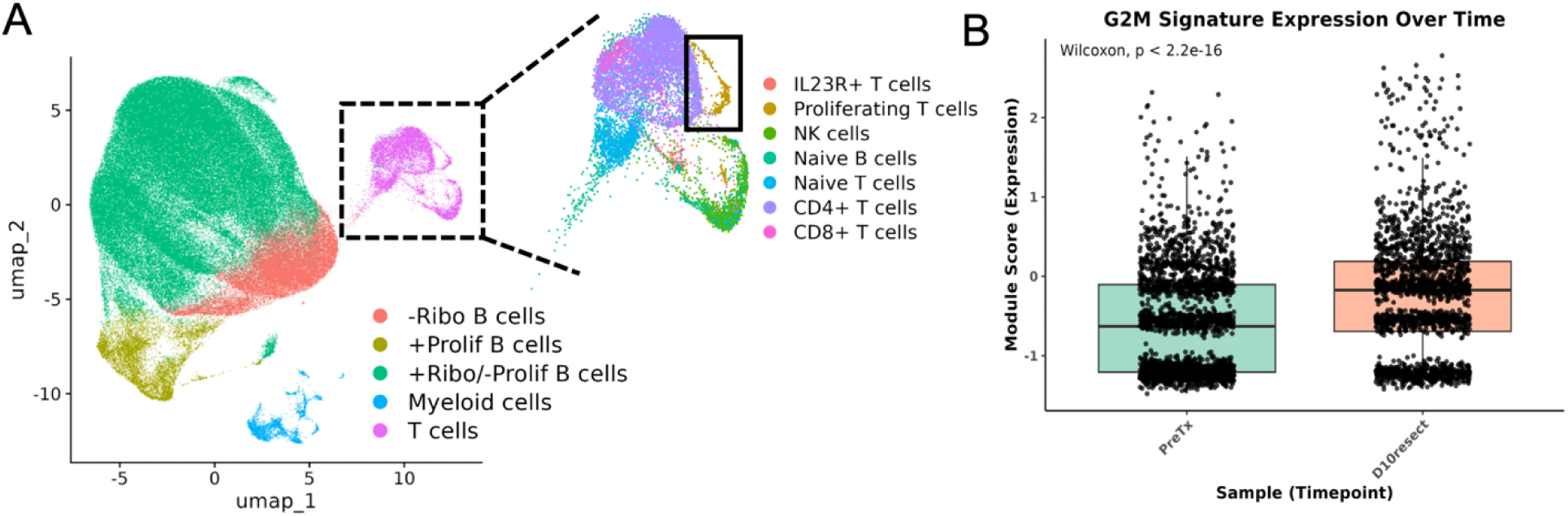
VSV induces a functionally intact anti-viral T-cell response despite a disrupted lymph node architecture. **A)** UMAP overview of sub-clustering the non-malignant lymphocyte cells from all timepoints. Proliferating T cells are highlighted by the solid black box. **B)** Boxplot of G2M gene signature expression between all non-malignant lymphocytes of PreTx and day 10 resection samples. Plot shows increased expression in the post treatment sample is largely attributed to a single cell population.

A focus on the proliferating T-cell subpopulation revealed dynamic changes in abundance across PostTx timepoints, with a noticeable expansion between days 7 and 10 following VSV administration (**Supplemental Fig. S6D**). Gene set scoring confirmed that this subset exhibited higher expression of G2/M cell cycle genes relative to all other populations (**Supplemental Fig. S6E**). Additionally, the increase in the G2/M gene set expression on days 7 to 10 is attributable to a small subset of cells from the non-malignant lymphocytes, while the remaining cells maintained relatively stable expression over time (**Fig. 4B, Supplemental Fig. S6F**). These results align with the canonical trajectory of an anti-viral response in T cells over time, peaking 7 to 10 days PostTx, and supporting the presence of T-cell activation and mobilization consistent with a primary anti-viral response.^30^

Differential gene expression and pathway analysis on T cells between individual timepoints Pre- and PostTx further supported the presence of an anti-viral T-cell response. Day 3 samples revealed upregulation of *GZMA*, *CTSW*, *NKG7, CD160,* and *KLRF1* when compared to PreTx samples (**Fig. 5A**). Such genes are highly implicated in T- and NK-cell activation and cytotoxicity.^31^ *IL12RB2* and *IL2RB,* induced upon antigen activation, along with the transcription factor *ETS1* required for T-cell activation, were similarly upregulated in the day 3 samples, as was *LTB*, a regulator of T-cell anti-viral immunity.^32,33,34^ Intriguingly, *PDCD4* also showed upregulation on day 3. While its expression is typically downregulated upon T-cell activation, upregulation has been shown to occur on virus-specific T cells prior to clonal expansion in both acute and persistent viral infections.^35,36^

**Figure 5.**
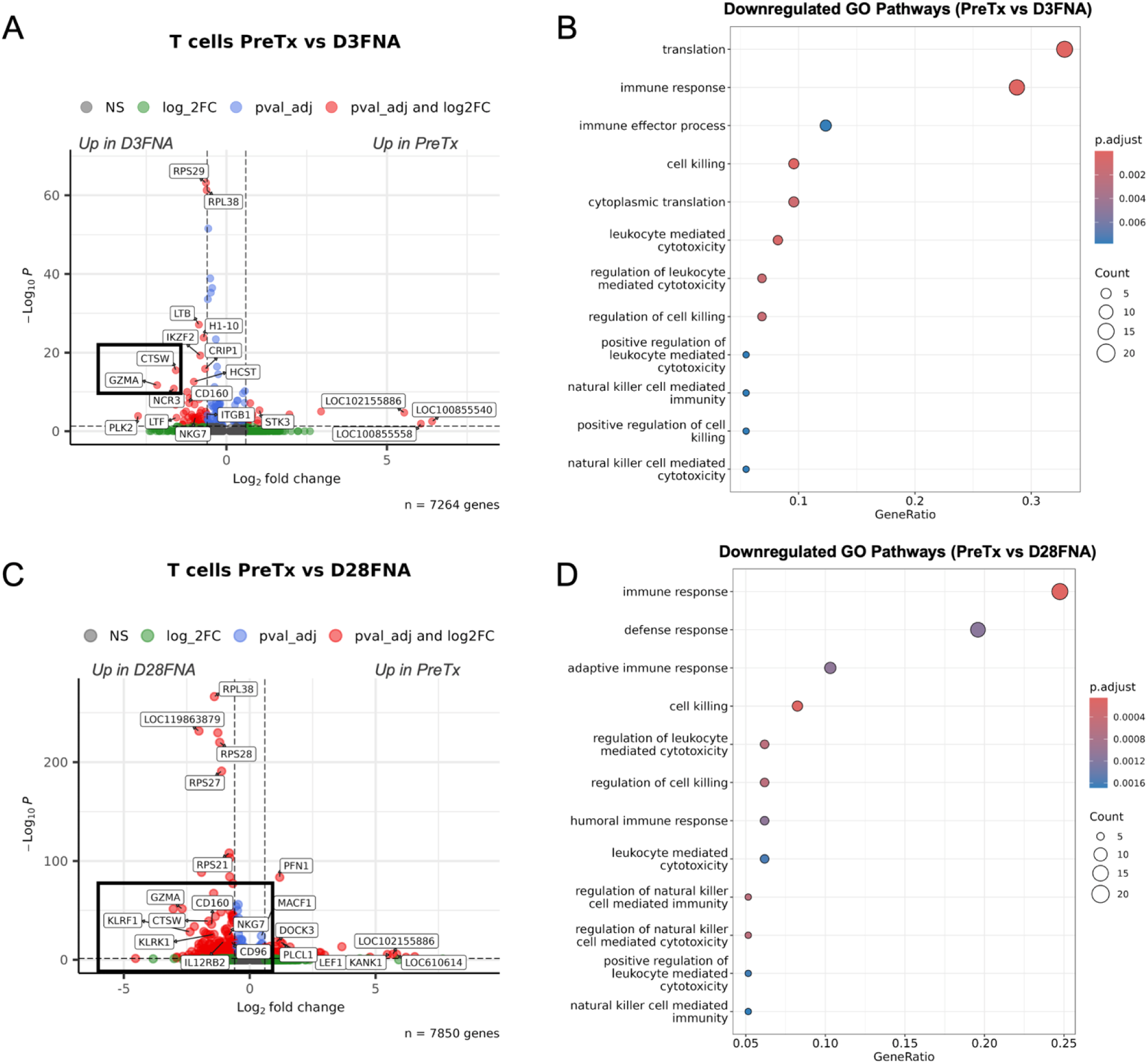
T-cell DGE and pathway analyses validate presence of a persistent anti-viral immune response. **A)** Volcano plot of differentially expressed genes between T cells of PreTx and day 3 samples. **B)** Enriched GO pathways of genes downregulated PreTx (*up on day 3*) from A. **C)** Volcano plot of differentially expressed genes between PreTx T cells and day 28 T cells. **D)** Enriched GO pathways of genes downregulated PreTx (*up on day 28*) from C.

Pathway analysis of genes upregulated on day 3 confirmed enrichment for key immune effector processes, including cell killing, immune effector process, leukocyte-mediated cytotoxicity, and NK cell-mediated immunity (**Fig. 5B**). This anti-viral response remained detectable at subsequent timepoints. While day 7 DEGs were dominated by ribosomal and cell cycle-associated genes (*MKI67, CENPF, CCNB2, CCNA2, ATP5F1E, RPS27, RPS29*) which limited enrichment of anti-viral pathways, cytotoxicity markers (*CTSW, CD52, GZMA*) still displayed concurrent upregulation, suggesting a transitional proliferative phase still consistent with expansion of activated T cells (**Supplemental Fig. S7A,B**).^30^ By day 10, pro-inflammatory cytokines showed downregulation while cytotoxic markers remained elevated, indicating progression toward an effector phenotype (**Supplemental Fig. S7C-E**). At day 28, T cells continued to exhibit enrichment for adaptive and humoral immune responses, cell killing regulation, and NK and leukocyte mediated cytotoxicity compared to PreTx, supporting sustained immune activation (**Fig. 5C,D**). Such findings collectively indicate a persistent and dynamic anti-viral immune response in the non-malignant compartment of the tumor, supporting the hypothesis that VSV induces a transcriptional change in these cells in response to the oncolytic pressure of tumor cell infection.

### Clonal B-cell subpopulations and non-malignant T lymphocytes occupy spatially segregated niches

Given the coexistence of presumptive clonal B-cell subpopulations and T cells implicated in an anti-viral immune response, we next sought to answer whether these populations displayed any spatial localization throughout the lymph node. Spatial transcriptomics of the day 10 resection yielded strong sequencing depth, but spot-based data resolution limited annotation of cell types by standard clustering and marker genes alone (**Supplemental Fig. S8**). Cell type deconvolution using the day 10 scRNAseq reference resolved this to reveal distinctive spatial niches in the tissue, with T cells and myeloid cells localizing to or near the lymph node trabeculae (enriched for fibrous stroma) and B-cell subpopulations occupying distinct regions throughout the lymph node (**Fig. 6A, Supplemental Fig. S9**). Non-proliferating B cells with a functioning ribosomal program showed the most widespread distribution across the lymph node space, while proliferating B cells appeared more localized to a distinct region in the upper right corner of the tissue image and ribosomal depleted B cells occupying the lower right of the image (**Fig. 6A**). CD4+ and CD8+ T cells were diffusely distributed throughout the tissue, consistent with effacement of normal lymph node architecture, in which follicular and interfollicular structures are replaced by malignant cells to create a more homogeneous appearance (**Fig. 6A**). The remaining spatial niches were supported by spatially variable gene expression analysis, indicating that genes *COL1A1* and *IGFBP7* were of the most spatially variable (**Fig. 6B**). Such genes emphasized the loyalty of cells to respective connective tissue or dispersed lymph node organization. Ribosomal genes such as *RPL37A* also exhibited strong spatial variability, however this likely reflects variation in RNA content captured across the tissue as opposed to robust spatial patterns of cellular organization.

**Figure 6.**
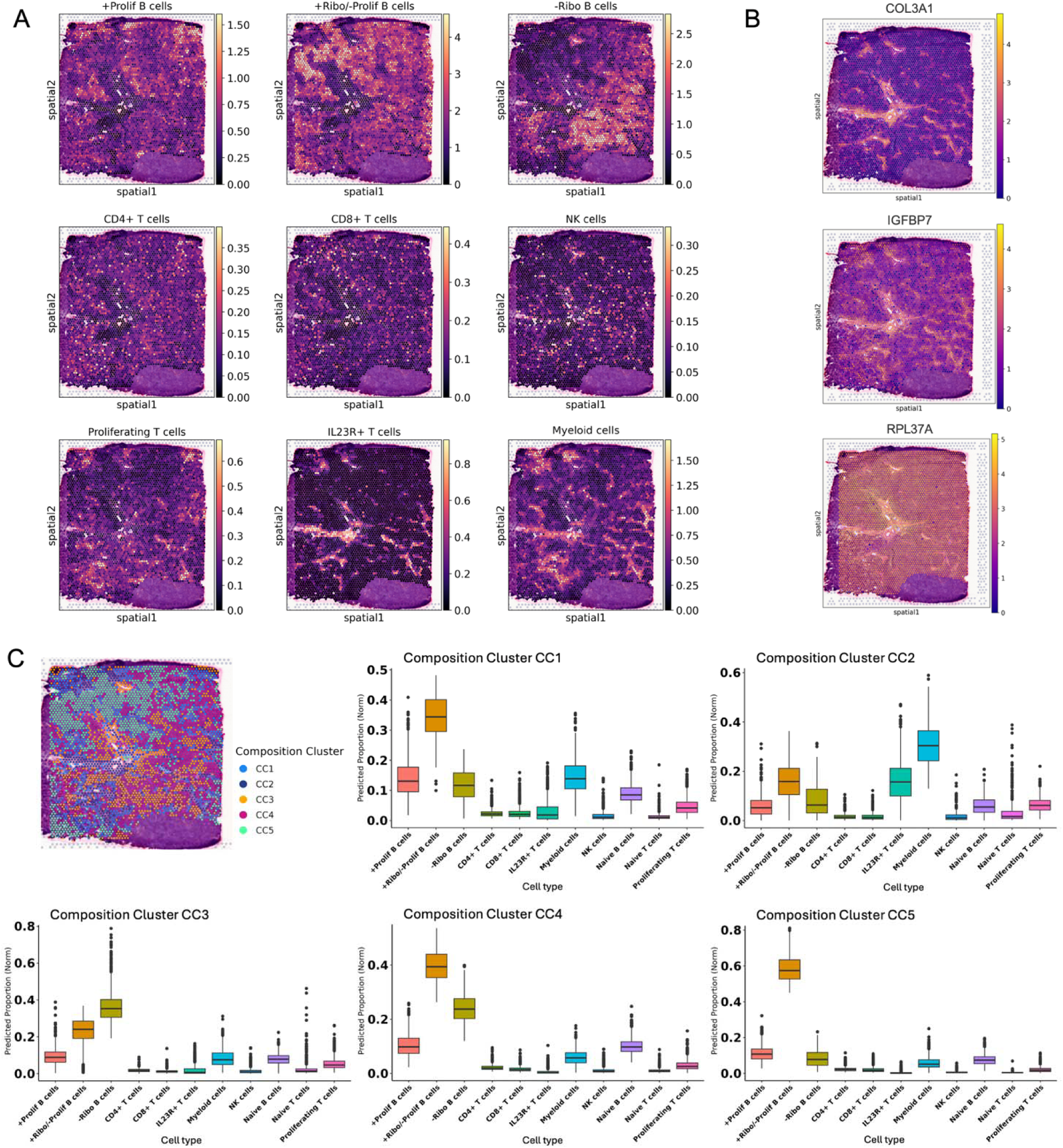
Clonal B-cell subpopulations and non-malignant T lymphocytes occupy spatially segregated niches. **A)** Spatial plots of cell2location predicted cell type proportions per 50 µm spot, using single-cell RNAseq data from the day 10 resection as a reference. **B)** Spatial plots of 3 top spatially variable genes present in the tumor. **C)** Spatial plot and associated boxplots showing results of composition clustering. Boxplots display cell type proportion profiles of spots within each composition cluster.

Cell composition-based clustering grouped spots into 5 niches based on their cell type prediction profiles (**Fig. 6C**), further revealing discrete spatial organization within the tumor. Composition clusters 1 and 2 represent higher enrichment of IL23+ T cells and myeloid cells, in contrast to clusters 1, 4, and 5 representing spots dominated by B cells (**Fig. 6C, Supplemental Fig. S10**). Not only do these results accurately reflect the previously established spatial architecture of the tumor, but a closer inspection of cluster cell type profiles reveals that de-enrichment of Ribo+ B cells coincides with an enrichment of Ribo-B cells, particularly in composition cluster 3 (**Fig. 6c**, **Supplemental Fig. S10**). Such spatial orientation suggests potential competitive dynamics among B-cell subpopulations and provides additional evidence to the transcriptional diversity present in the tumor.

## Discussion

Diffuse B-cell lymphomas are the most common canine hematological malignancies, but the detailed molecular processes driving subtype stratification remain unclear. For canine MZL in particular, a poor understanding of its biology impedes development of effective strategies to address complex mechanisms of therapy resistance. Oncolytic virotherapy shows promise to improve treatment response and durable remissions, but its intratumoral effects have not been fully characterized at the transcriptional level. To address these gaps, we leveraged the depth supplied by a longitudinal, single-subject design of combined scRNAseq and spatial transcriptomics approaches to construct a high-resolution molecular framework of canine MZL cellular composition and therapeutic response.

Our results indicate a complex intratumoral heterogeneity, consisting of three transcriptionally distinct B-cell subpopulations whose displayed stability under oncolytic pressure and robust spatial organization implicates an intrinsic, heterogeneous tumor architecture. Further analysis of these MZL B cells identified copy number variations and a collective downregulation of the quiescence program. Within the T-cell compartment of the tumor, we detected an anti-viral immune response that persisted throughout the treatment time course. Collectively our findings present a unique assessment of the temporal transcriptional dynamics of a canine MZL, offering insight to comparative molecular signatures and viral-specific changes present in the tumor.

Despite the lack of VSV-induced transcriptional changes in the malignant B cells of this tumor, our identification of three unique and longitudinally stable B-cell subpopulations both before and after treatment supports that intratumoral heterogeneity of canine MZL may be a primary obstacle to effective immunotherapeutic approaches. Each subpopulation displayed a distinct transcriptional profile, indicating a complex cellular plasticity within the tumor that VSV monotherapy did not appear to fully abrogate. The downregulated ribosomal signature that delineates one of the three B-cell subgroups has previously been implicated in human diffuse B-cell lymphoma; it is specifically reported in human diffuse large B-cell lymphoma (DLBCL) and in instances where tumor cells exist in a state of hypoxia.^37,38^ However, this signature has not yet been reported in canine MZL, and its contribution to tumor progression remains largely unexplored. Its identification here illustrates the utility of high-resolution single-cell profiling for uncovering subtype-specific biology, amenable to future multi-subject investigation and validation.

The stability of each B-cell subpopulation, despite the presence of oncolytic pressure, further suggests that the tumor’s clonal evolution does not progress on a weeks-long timescale. The persistence of the populations over the treatment time course instead indicates a clonal architecture resistant to VSV-driven adaptations. Our inability to detect VSV transcripts in the single-cell data was likely due to the relatively low abundance of viral transcripts, as robust copies were detectable on day 3 with the higher sensitivity targeted qRT-PCR assay. Alternatively, the administered viral dose may have been inadequate to achieve direct, widespread oncolytic lysis in this dog (one fifth of the maximal tolerated dose).^8^ It warrants consideration that technical biopsy variation and subsequent differences in sample quality constrained direct comparisons across timepoints. While reported biological results showed consistency across the treatment time course, implications from possible batch effects must be acknowledged when interpreting such results. Nevertheless, the observed clonal stability underscores an imperative need for multidimensional therapeutic strategies capable of simultaneously targeting the multiple molecular mechanisms operating within the tumor.

The diversification of CNVs predicted within the -Ribo B-cell subpopulation, some overlapping with those predicted in the +Ribo/-Prolif B cells, further emphasizes a complex tumor heterogeneity and evolution. Plausible biological explanations include a single clonal population existing in several transcriptional states, or a replicating tumor population differentiating into ribosomal high and low groups alongside non-malignant B cells. Additional work to determine how certain CNVs, such as loss on CFA20 with its concomitant loss of *KLF2*, may be driving malignancy would help clarify the similarities and differences between mechanisms contributing to canine and human marginal zone lymphomagenesis. The single-subject nature of this study is an inherent constraint on population-level generalizability, and an expanded cohort will be essential to validate conclusions regarding CNV-driven tumor pathogenesis and specific molecular drivers of the disease. However, the depth of molecular characterization presented here provides a strong preliminary foundation for such future multi-subject studies.

We reported downregulation of the transcription factor *KLF2* in the serial B cells that is consistent with previous reports in human MZL. *KLF2* normally functions as a tumor suppressor in other cancers but appears frequently mutated in human SMZL, resulting in inactivation and subsequent deregulation of critical pathways with oncogenic repercussions.^39,40,41^ *BACH2* and *RIPOR2,* also downregulated, are transcription factors implicated in the support and maintenance of a quiescent cell state in lymphocytes.^42,43^ Their decreased expression is consistent with a shift towards activation and proliferation in the serial B cells. Overall, our findings provide strong preliminary evidence that, despite developing in different anatomical locations, this canine NMZL recapitulated oncogenic mechanisms seen in its splenic human counterpart. This potentially positions the canine disease as a practical comparative model with unrealized translation relevance, requiring clearer subtyping framework to more precisely characterize how such conserved mechanisms drive tumor progression in both species.

We also identified proliferating T cells PostTx and differential expression of cytotoxic markers and related pathways to support that treatment of canine MZL with VSV is associated with an anti-viral immune response. The priming and activation of non-malignant lymphocytes, coinciding with the peak detection of viral transcripts, specifically suggests that the cells at minimum encountered the virus and subsequently elicited an immune response that persisted through day 28. This result is also consistent with prior observations of activated T-cell infiltration in canine MZL.^3^ Such therapeutic response durability highlights the importance of longitudinal monitoring to capture the temporal dynamics of both tumor and host immune system relative to immunotherapy, and it also supports the continued development of VSV-based combination strategies for treatment of canine lymphoma.

Finally, we report a spatial transcriptomics analysis of this MZL tumor complementary to our single-cell findings, highlighting a complex lymph node composition, both homogenously distributed and yet disorganized in nature. We observed localization of cell types to unique niches within the tissue, and composition-based clustering emphasizes a relative enrichment and de-enrichment of non-malignant populations and B-cell groups across lymph node space. The unique and seemingly segregated localization of each B-cell subpopulation is particularly notable, suggesting a potentially competitive tumor environment in which each group prioritizes distinct spatial niches that may support divergent functional states or varying immunotherapeutic responses. Taken together with our single-cell findings, we showcase the utility of a multi-modal transcriptomics approach to investigate the intratumoral heterogeneity of canine MZL, resolving the transcriptional complexity within the tumor and its implications on molecular phenotype and immunotherapy response.

## Acknowledgements

This study was funded by grant U01 CA272257 from the National Cancer Institute (NCI) of the National Institutes of Health (NIH) of the US Public Health Service, and by the Animal Cancer Care and Research Program (ACCRP) and its generous donors. This work was also supported in part by the NCI Comprehensive Cancer Center Support Grant P30 CA77598 utilizing the following Masonic Cancer Center, University of Minnesota shared resource(s): Comparative Pathology, Cancer Informatics, and infrastructure support, by the NIH’s National Center for Advancing Translational Sciences (NCATS), grant UM1 TR004405, and by resources and staff at the University of Minnesota University Imaging Centers (RRID: SCR_020997), and the University of Minnesota Genomics Center (RRID: SCR_012413). JFM is supported in part by the Alvin and June Perlman Endowed Chair in Animal Oncology. AET was supported in part by a fellowship from the NCI (U01CA272257-S1). The content of this manuscript is solely the responsibility of the authors and does not necessarily represent the official views of any of the funders. The authors thank the University Imaging Center (UIC) for assistance with imaging of spatial transcriptomics slides; the University of Minnesota Genomics Center (UMGC) for single cell GEM capture, library preparation, quality control, and sequencing; the Minnesota Supercomputing Institute (MSI) for sequencing data processing support and storage; the Clinical and Translational Science Institute (CTSI), specifically Colleen Forster for training on 10X Visium Fresh Frozen tissue protocols and for preparing Tissue Optimization slides; the Department of Ophthalmology and Visual Neurosciences Histology Core Laboratory, specifically Heidi Roehrich for sectioning and placing fresh frozen tissues on Visium slides; the Comparative Pathology Shared Resource (CPSR), specifically Katalin Kovacs and Paula Overn who assisted with the preparation of samples for histopathology analyses; the College of Veterinary Medicine’s Clinical Investigation Center (CIC), as well as the Lewis Small Animal Hospital’s Oncology and Surgery Services for clinical study resource support, sample collection, diagnostics, and patient care.

## Author’s contributions

SN developed the VSV therapeutic platform and, with JFM, secured NIH funding. ALS, AET, AB, DAV, SN, and JFM contributed to conceptualization. JFM and AET oversaw experimental design. ALS provided custom bioinformatics workflows and supervised data analysis and visualization. AET, LEB, and SN performed sample processing and spatial library preparations. DMS and KDD performed immunophenotyping and sub-diagnosis. GEW and MM performed bioinformatics data curation, investigation, and analysis. VP processed samples and completed pharmacokinetic analysis. AB and MH contributed to dog recruitment, and with CY, RA, RD, CF, and AK, provided clinical management, treatment, and sample collection. KR, AW, SP, and MB provided clinical resource support, assisting with clinical management and performed sample collection. CF contributed to project administration, methodology, and clinical supervision. GEW primarily wrote the manuscript with contributions from several co-authors. All authors read and approved the final version.

## References

1. Hughes, K. L. et al. Diffuse Small B-Cell Lymphoma: A High-Grade Malignancy. Vet. Pathol. 58, 912–922 (2021).

2. Ito, D., Frantz, A. M. & Modiano, J. F. Canine lymphoma as a comparative model for human non-Hodgkin lymphoma: recent progress and applications. Vet. Immunol. Immunopathol. 159, 192–201 (2014).

3. Frantz, A. M. et al. Molecular Profiling Reveals Prognostically Significant Subtypes of Canine Lymphoma. Vet. Pathol. 50, 693–703 (2013).

4. Laurent, C. & Bertoni, F. The biology of marginal zone lymphoma subtypes: challenge and relevance of classification. Blood 147, 105–114 (2026).

5. Cheah, C. Y., Zucca, E., Rossi, D. & Habermann, T. M. Marginal zone lymphoma: present status and future perspectives. Haematologica 107, 35–43 (2022).

6. Childress, M. O., Ramos-Vara, J. A. & Ruple, A. Retrospective analysis of factors affecting clinical outcome following CHOP-based chemotherapy in dogs with primary nodal diffuse large B-cell lymphoma. Vet. Comp. Oncol. 16, (2018).

7. Flory, A. B. et al. Evaluation of factors associated with second remission in dogs with lymphoma undergoing retreatment with a cyclophosphamide, doxorubicin, vincristine, and prednisone chemotherapy protocol: 95 cases (2000–2007). J. Am. Vet. Med. Assoc. 238, 501–506 (2011).

8. Naik, S. et al. Comparative Oncology Evaluation of Intravenous Recombinant Oncolytic Vesicular Stomatitis Virus Therapy in Spontaneous Canine Cancer. Mol. Cancer Ther. 17, 316–326 (2018).

9. Hemminki, O., Dos Santos, J. M. & Hemminki, A. Oncolytic viruses for cancer immunotherapy. J. Hematol. Oncol.J Hematol Oncol 13, 84 (2020).

10. Yeung, K. T., et al. LBA2 Rates of pathologic complete response after VV1 and cemiplimab (alone and in treatment sequence-strategy) in the neoadjuvant setting: Results from the I-SPY 2.2 trial. ESMO Open 10, (2025).

11. Razmara, A. M. et al. Comparative transcriptomic analysis of tumor-infiltrating canine natural killer cells and candidate biomarkers from first-in-dog NK immunotherapy trials. Front. Immunol. 16, (2025).

12. Dittrich, K. et al. Baseline tumor gene expression signatures correlate with chemoimmunotherapy treatment responsiveness in canine B cell lymphoma. PloS One 18, e0290428 (2023).

13. Li, T. et al. Integrated single-cell transcriptome and TCR profiles of hepatocellular carcinoma highlight the convergence on interferon signaling during immunotherapy. J. Immunother. Cancer 12, e010534 (2024).

14. Cui, X., Liu, S., Song, H., Xu, J. & Sun, Y. Single-cell and spatial transcriptomic analyses revealing tumor microenvironment remodeling after neoadjuvant chemoimmunotherapy in non-small cell lung cancer. Mol. Cancer 24, 111 (2025).

15. Cambien, B. et al. Identification of oncolytic vaccinia restriction factors in canine high-grade mammary tumor cells using single-cell transcriptomics. PLOS Pathog. 16, e1008660 (2020).

16. Chen, C. et al. Single-cell panleukemia signatures of HSPC-like blasts predict drug response and clinical outcome. Blood 145, 2685–2700 (2025).

17. Iacobucci, I. et al. Multipotent lineage potential in B cell acute lymphoblastic leukemia is associated with distinct cellular origins and clinical features. *Nat*. Cancer 6, 1242–1262 (2025).

18. Lou, N., et al. Single-cell sequencing and spatial transcriptomics reveal FAS+ T cell and autophagy-related signatures predicting chemoimmunotherapy response in diffuse large B-cell lymphoma patients. Sci. China Life Sci. 68, 2316–2331 (2025).

19. Sklavenitis-Pistofidis, R. et al. Single-cell RNA sequencing defines distinct disease subtypes and reveals hypo-responsiveness to interferon in asymptomatic Waldenstrom’s Macroglobulinemia. Nat. Commun. 16, 1480 (2025).

20. Seelig, D. M. et al. Canine T-zone lymphoma: unique immunophenotypic features, outcome, and population characteristics. J. Vet. Intern. Med. 28, 878–886 (2014).

21. Valli, V. E. et al. Classification of canine malignant lymphomas according to the World Health Organization criteria. Vet. Pathol. 48, 198–211 (2011).

22. Flood-Knapik, K. E. et al. Clinical, histopathological and immunohistochemical characterization of canine indolent lymphoma. Vet. Comp. Oncol. 11, 272–286 (2013).

23. Dyer, S. C. et al. Ensembl 2025. Nucleic Acids Res. 53, D948–D957 (2025).

24. Goldfarb, T. et al. NCBI RefSeq: reference sequence standards through 25 years of curation and annotation. Nucleic Acids Res. 53, D243–D257 (2025).

25. Tirosh, I. et al. Dissecting the multicellular ecosystem of metastatic melanoma by single-cell RNA-seq. Science 352, 189–196 (2016).

26. Scott, M. C. et al. Comparative Transcriptome Analysis Quantifies Immune Cell Transcript Levels, Metastatic Progression, and Survival in Osteosarcoma. Cancer Res. 78, 326–337 (2018).

27. Aricò, A. et al. Array-Based Comparative Genomic Hybridization Analysis Reveals Chromosomal Copy Number Aberrations Associated with Clinical Outcome in Canine Diffuse Large B-Cell Lymphoma. PLoS ONE 9, e111817 (2014).

28. Thomas, R. et al. Refining tumor-associated aneuploidy through ‘genomic recoding’ of recurrent DNA copy number aberrations in 150 canine non-Hodgkin lymphomas. Leuk. Lymphoma 52, 1321–1335 (2011).

29. Campos-Martín, Y. et al. Clinical and diagnostic relevance of *NOTCH2* -and *KLF2* - mutations in splenic marginal zone lymphoma. Haematologica 102, e310–e312 (2017).

30. Sun, L., Su, Y., Jiao, A., Wang, X. & Zhang, B. T cells in health and disease. Signal Transduct. Target. Ther. 8, 235 (2023).

31. Stoeckle, C. et al. Cathepsin W expressed exclusively in CD8+ T cells and NK cells, is secreted during target cell killing but is not essential for cytotoxicity in human CTLs. Exp. Hematol. 37, 266–275 (2009).

32. Szabo, S. J., Dighe, A. S., Gubler, U. & Murphy, K. M. Regulation of the Interleukin (IL)-12R b2 Subunit Expression in Developing T Helper 1 (Th1) and Th2 Cells.

33. Berger, D. P. et al. Lymphotoxin-b-Deficient Mice Show Defective Antiviral Immunity.

34. Szabo, S. J., Dighe, A. S., Gubler, U. & Murphy, K. M. Regulation of the Interleukin (IL)-12R b2 Subunit Expression in Developing T Helper 1 (Th1) and Th2 Cells.

35. Ahn, E. et al. Role of PD-1 during effector CD8 T cell differentiation. Proc. Natl. Acad. Sci. 115, 4749–4754 (2018).

36. Schönrich, G. & Raftery, M. J. The PD-1/PD-L1 Axis and Virus Infections: A Delicate Balance. Front. Cell. Infect. Microbiol. 9, 207 (2019).

37. Wu, W. et al. Prognostic Significance of Ribosome-related Genes Signature in Diffuse Large B Cell Lymphoma. J. Cancer 14, 403–416 (2023).

38. Duś-Szachniewicz, K., Gdesz-Birula, K., Zduniak, K. & Wiśniewski, J. R. Proteomic-Based Analysis of Hypoxia- and Physioxia-Responsive Proteins and Pathways in Diffuse Large B-Cell Lymphoma. Cells 10, 2025 (2021).

39. Wang, C., Li, L., Duan, Q., Wang, Q. & Chen, J. Krüppel-like factor 2 suppresses human gastric tumorigenesis through inhibiting PTEN/AKT signaling. Oncotarget 8, 100358–100370 (2017).

40. Li, Y.-Z. et al. KLF2 is a clinical diagnostic and treatment biomarker of breast cancer. Front. Cell Dev. Biol. 11, 1182123 (2023).

41. Clipson, A. et al. KLF2 mutation is the most frequent somatic change in splenic marginal zone lymphoma and identifies a subset with distinct genotype. Leukemia 29, 1177–1185 (2015).

42. Grant, F. M. et al. BACH2 drives quiescence and maintenance of resting Treg cells to promote homeostasis and cancer immunosuppression. J. Exp. Med. 217, e20190711 (2020).

43. Lv, Z., Ding, Y., Cao, W., Wang, S. & Gao, K. Role of RHO family interacting cell polarization regulators (RIPORs) in health and disease: Recent advances and prospects. Int. J. Biol. Sci. 18, 800–808 (2022).

