## Supplementary Data for "Longitudinal single-cell and spatial transcriptomics reveals intratumor heterogeneity, therapeutic response, and comparative value of canine marginal zone lymphoma"

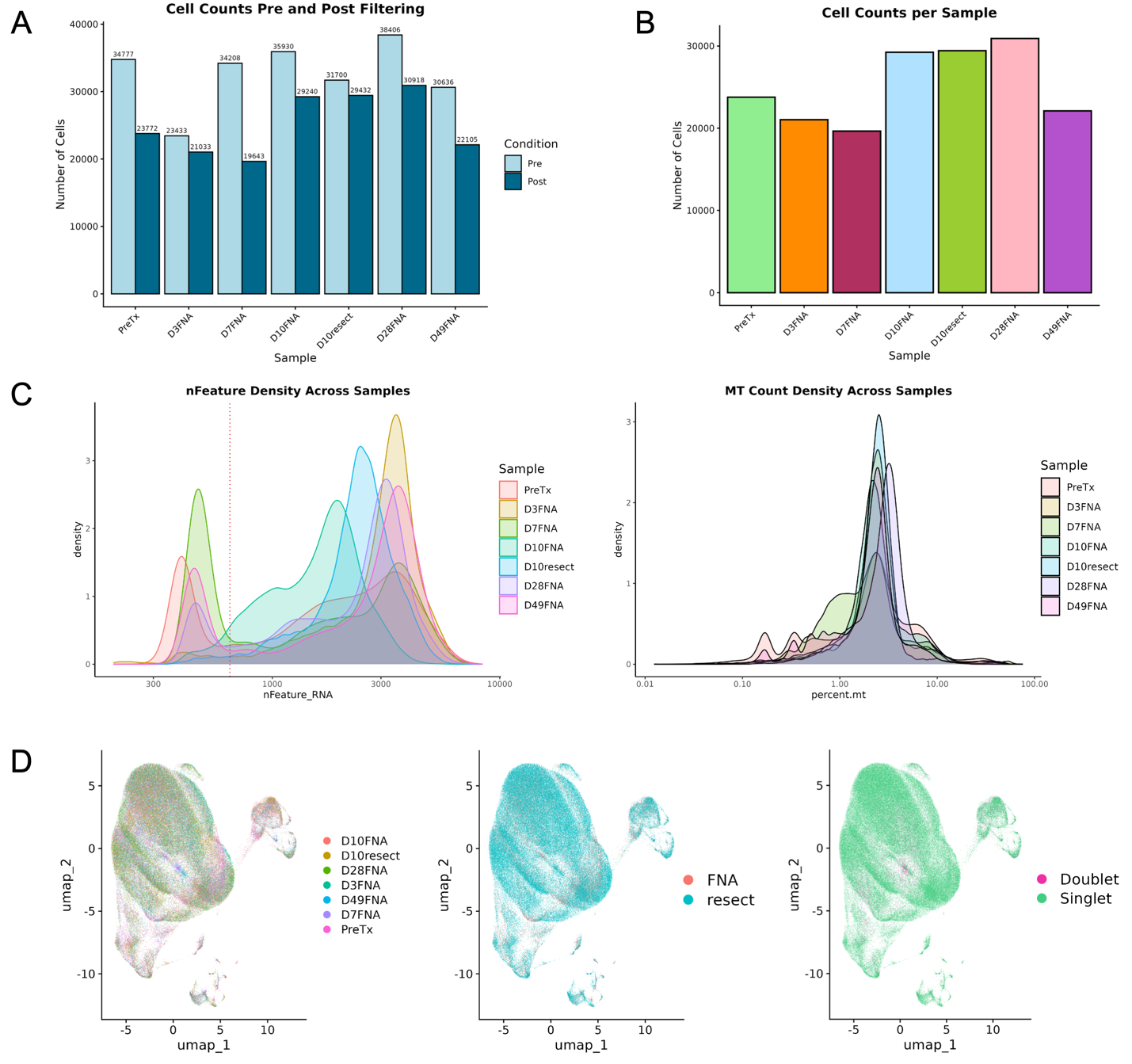

**Supplementary Figure 1. Quality control and cell filtering confirm robust sample integrity despite technical biopsy variability. A)** Serial scRNAseq cell counts pre- and post-quality control. **B)** Final cell counts across serial scRNAseq samples. **C)** Density plots of unique features (*left*) and mitochondrial gene expression (*right*) distributions for each timepoint. **D)** Aggregated UMAPs colored by sample, technical biopsy method, and doublet prediction.

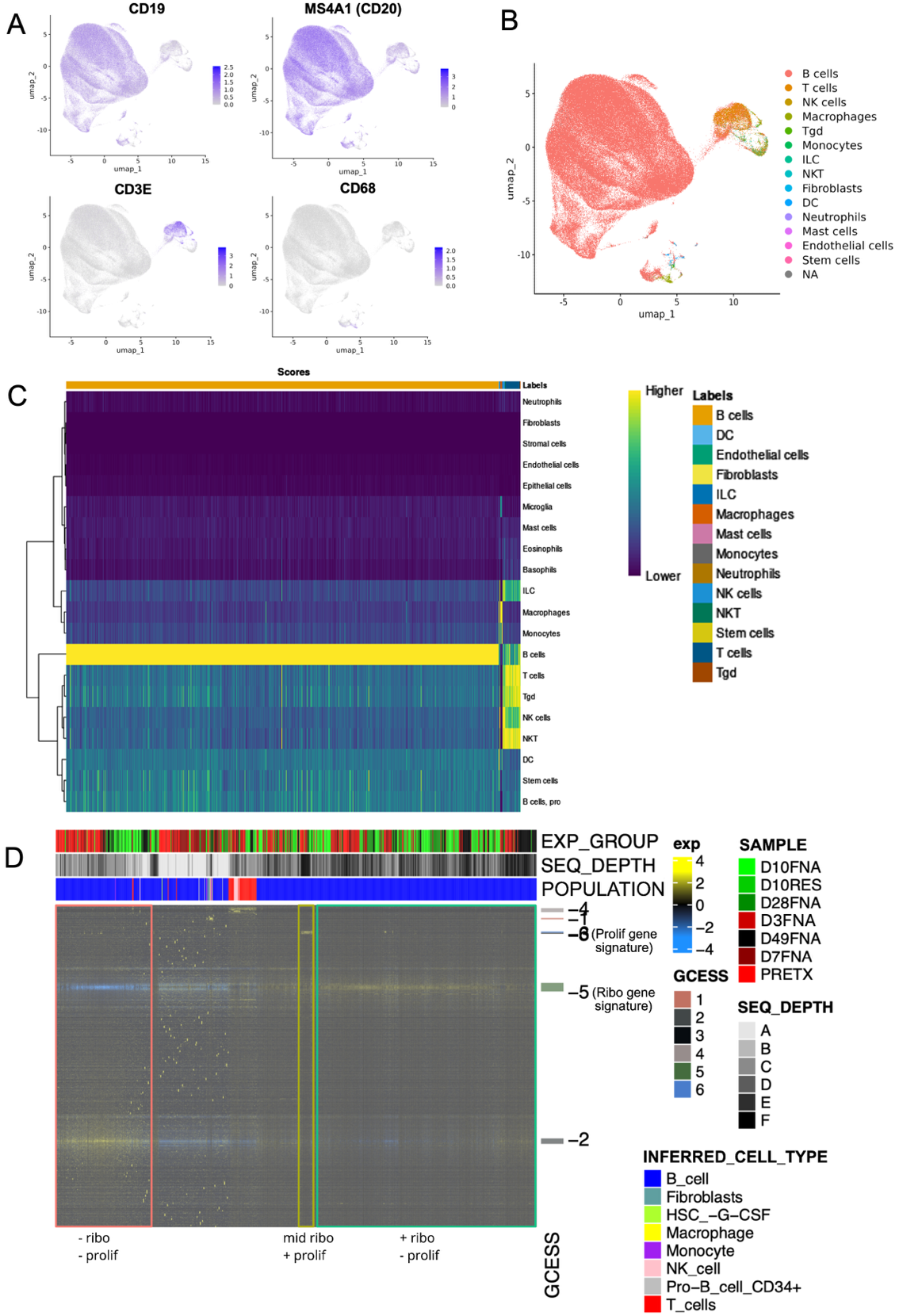

**Supplementary Figure 2. Automated annotation and GCESS analysis validate presence and stability of B-cell subpopulations over treatment time course. A)** Cell marker UMAPs for manual annotation. **B)** UMAP of SingleR automated cell type predictions using the ImmGen reference dataset. **C)** Heatmap of prediction scores for B. **D)** Heatmap of predicted expression clusters and cell types with GCESS. Unique B-cell subpopulations are indicated by the red, yellow, and green boxes.

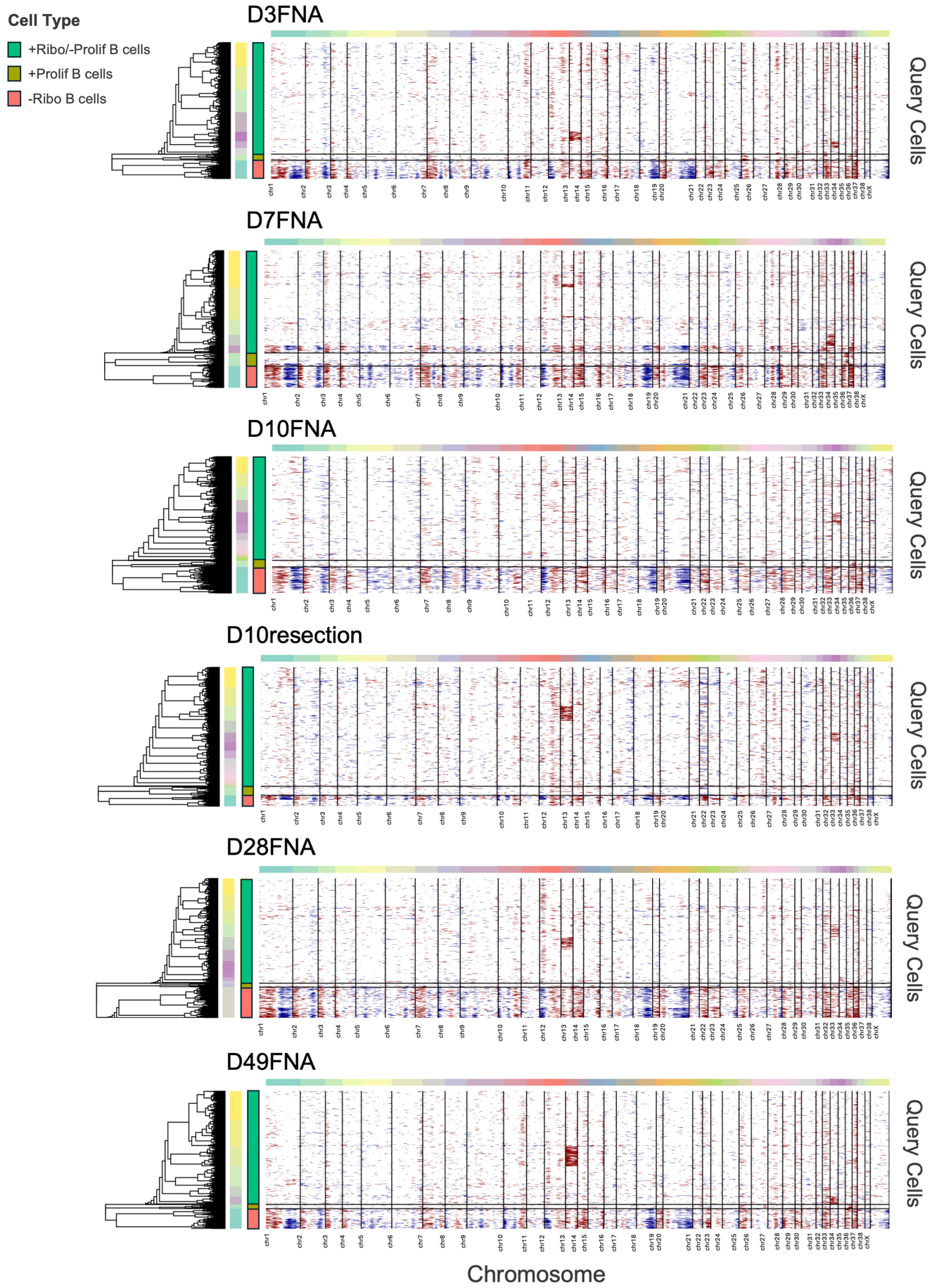
 **Supplementary Figure 3. Predicted copy number alterations are persistent across all timepoints.** Heatmaps of InferCNV copy number profile predictions for each timepoint.

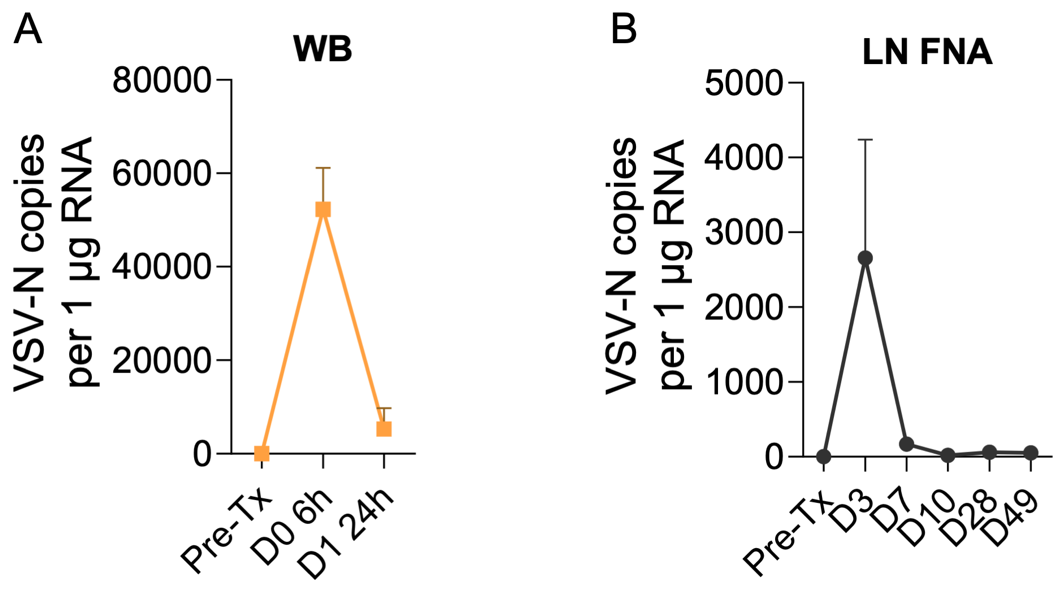

**Supplementary Figure 4. Evidence of systemic delivery and localized intratumoral replication of VSV.** Targeted qRT-PCR detection of VSV-N transcripts in **A)** whole blood at 6h and 24h following VSV treatment, and **B)** in serial lymph node FNA samples following VSV treatment.

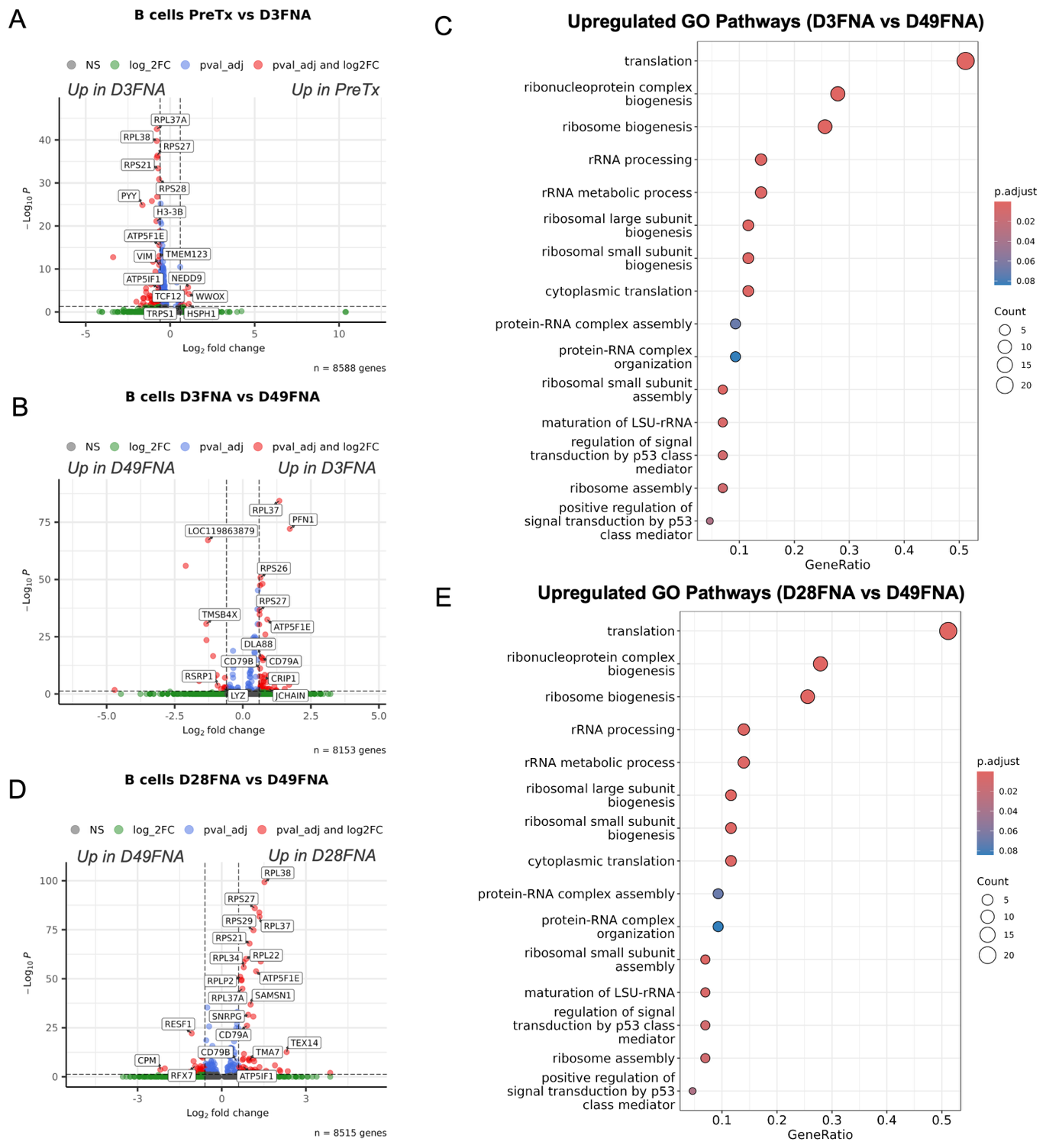

**Supplementary Figure 5. Oncolytic pressure from VSV treatment did not induce transcriptional changes in B cells over time. A)** Volcano plot of differentially expressed genes between PreTx B cells and day 3 B cells. **B)** Volcano plot of differentially expressed genes between day 3 B cells and day 49 B cells. **C)** Enriched GO pathways of genes upregulated on day 3 (*down on day 49*) from B. **D)** Volcano plot of differentially expressed genes between day 28 B cells and day 49 B cells. **E)** Enriched GO pathways of genes upregulated on day 28 (*down on day 49*) from D.

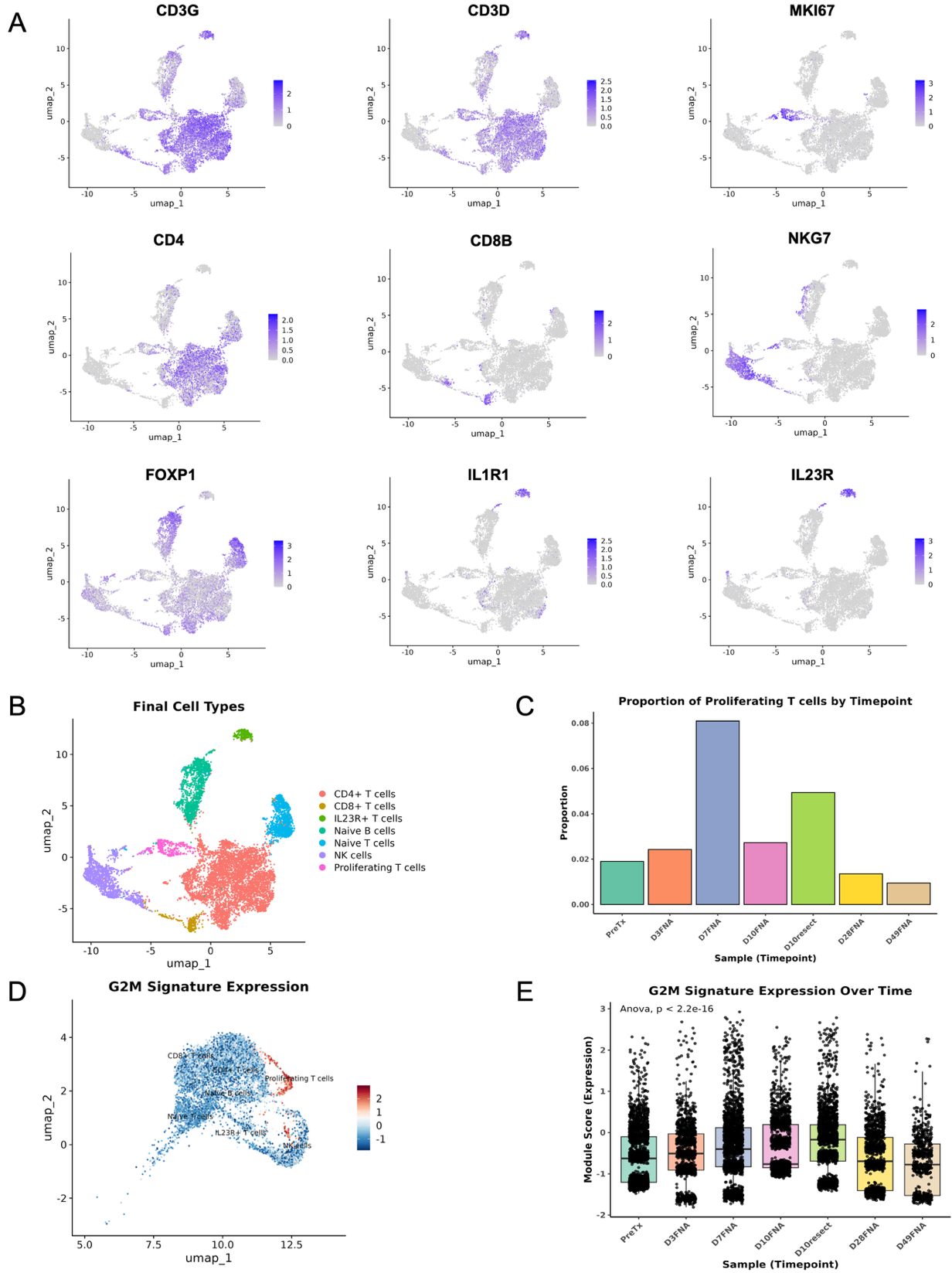

**Supplementary Figure 6. Proliferating T-cell subpopulation is present in varying proportions across all timepoints. A)** Cell type marker UMAPs for manual annotation. **B)** Final cell type annotations from T-cell subclustering. **C)** Proportion of proliferating T cells across serial samples. **D)** UMAP of G2M gene signature expression in the T cell population across all timepoints. **E)** Boxplot of G2M gene signature expression between all non-malignant lymphocytes, across all timepoints.

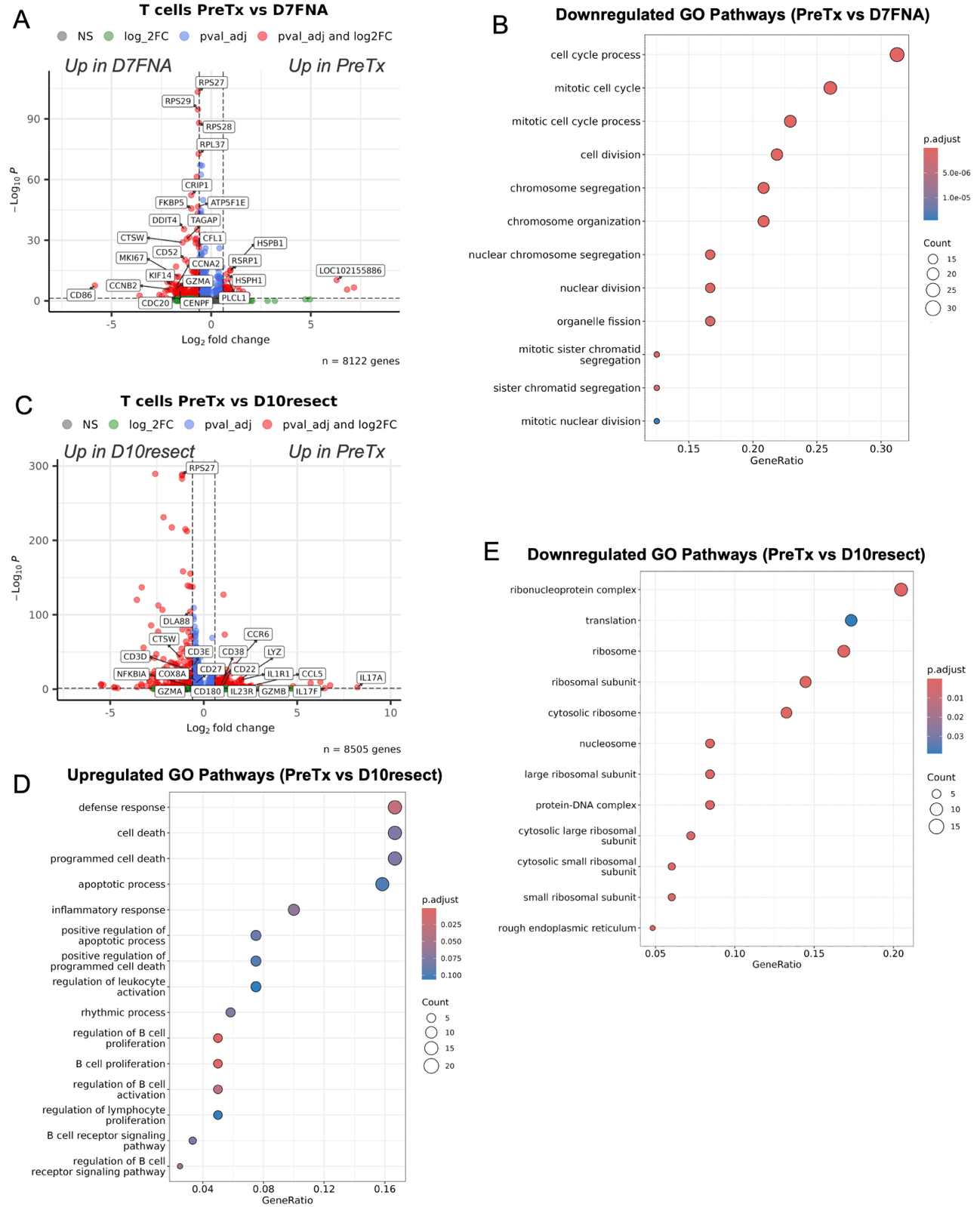

**Supplementary Figure 7. Anti-viral T-cell response persists across timepoints. A)** Volcano plot of differentially expressed genes between PreTx T cells and day 7 T cells. **B)** Enriched GO pathways of genes downregulated PreTx (*up on day 7*) from A. **C)** Volcano plot of differentially expressed genes between PreTx T cells and day 10 (resection) T cells. **D)** Enriched GO pathways of genes downregulated PreTx (*up on day 10 resection*) from C. **E)** Enriched GO pathways of genes upregulated PreTx (*down on day 10 resection*) from C.

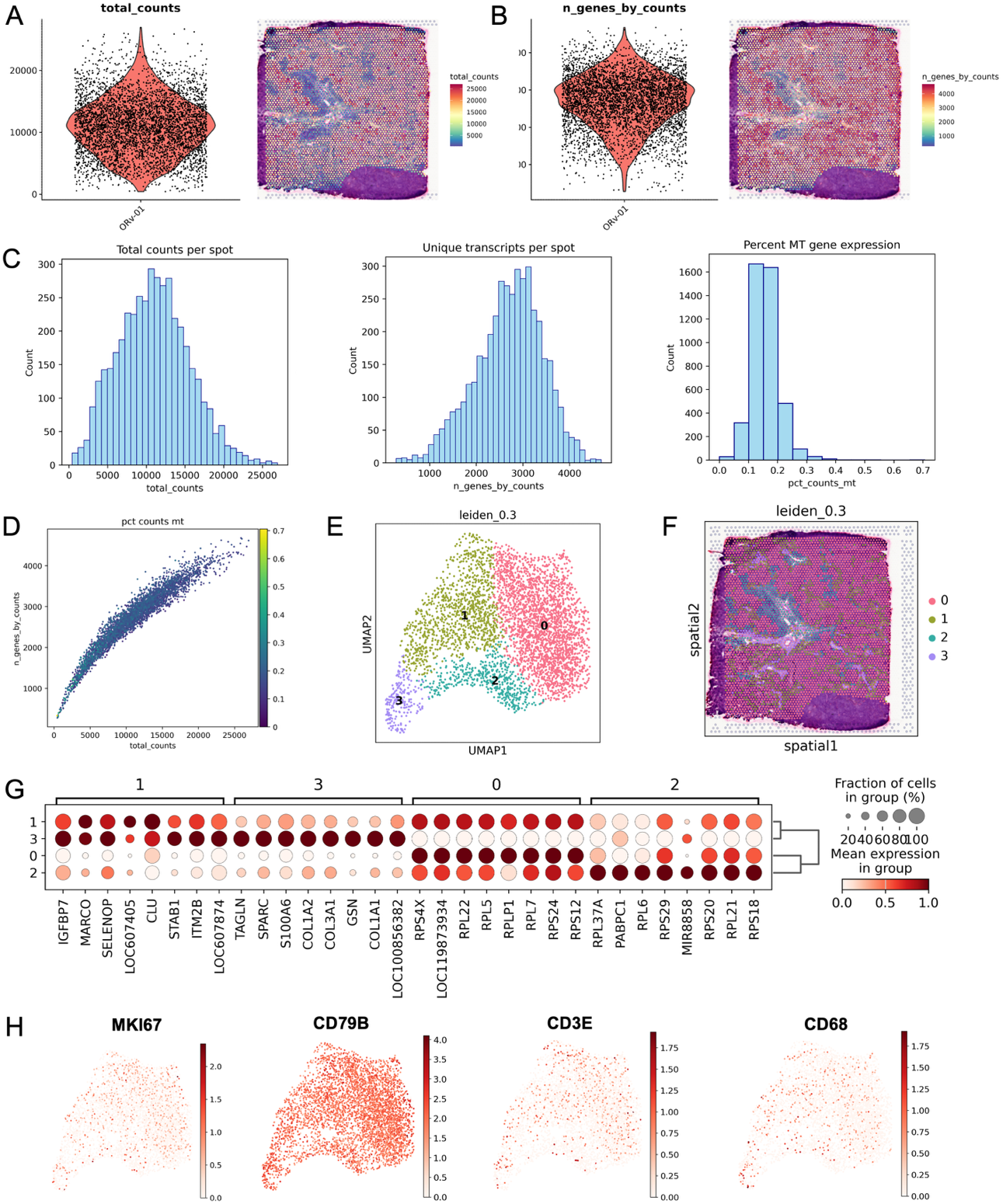

**Supplemental Figure 8. Spatial quality control and filtering confirms strong sequencing depth and minimal contaminant spots. A)** Violin and spatial plots of total counts in the spatial dataset. **B)** Violin and spatial plots of unique feature counts in the spatial dataset. **C)** Histograms of total counts, unique transcripts, and percent MT gene expression per spatial spot. **D)** Scatter plot of total counts by unique features per spatial spot. Color bar indicates % MT expression. **E)** UMAP of leiden clustering results. **F)** Spatial plot of clustering results from e. **G)** Dotplot of top 8 differentially expressed genes per leiden cluster. **H)** B-cell, T-cell, and myeloid marker gene plots.

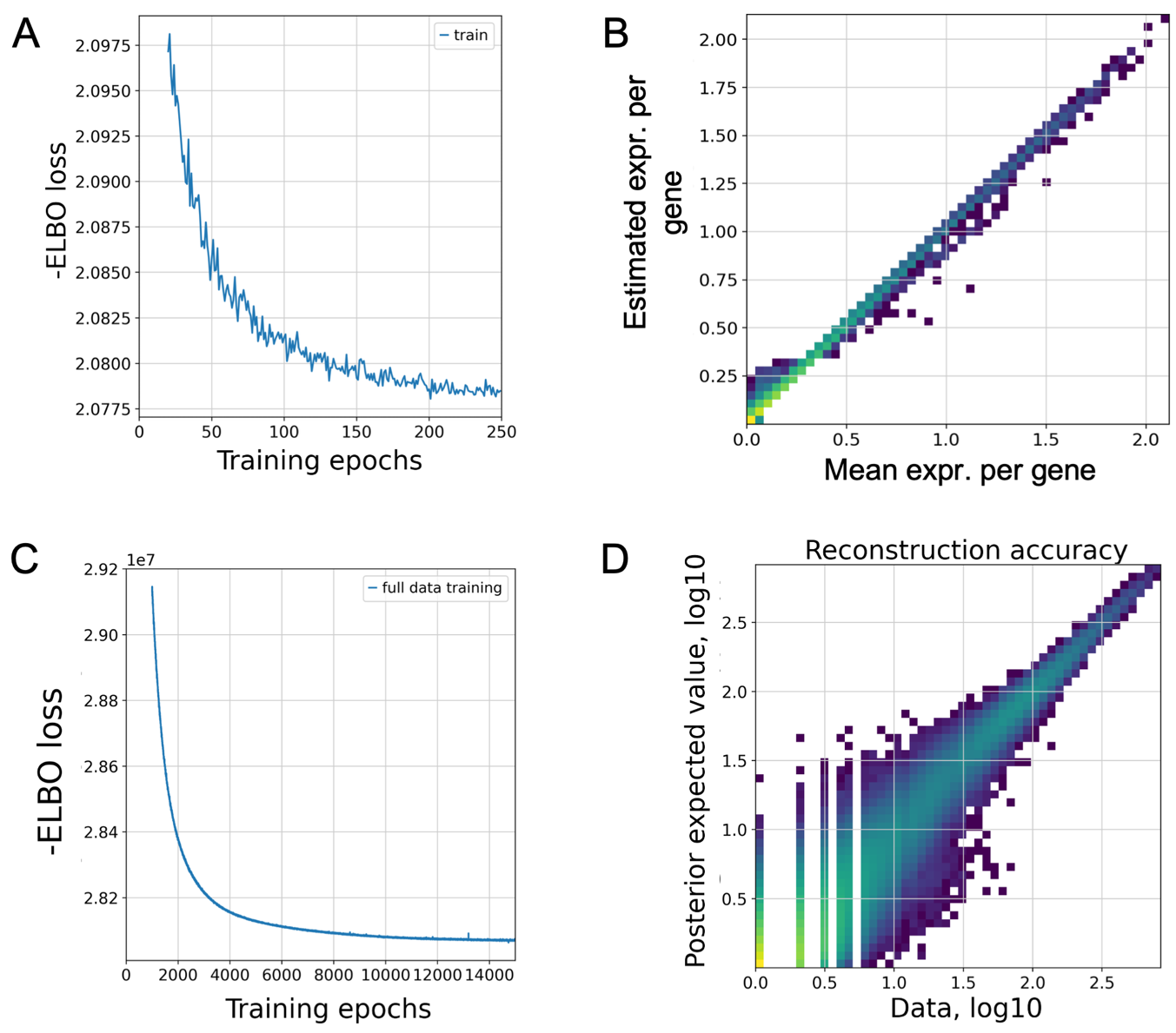

**Supplementary Figure 9. Cell2location deconvolution model shows minimal loss and high accuracy when applied to Visium v1 data. A)** ELBO loss across model training for part one of the c2l deconvolution algorithm. **B)** Scatter plot of model estimated expression signatures against mean expression per gene. **C)** ELBO loss across model training for part 2 of the c2l algorithm. **D)** Scatter plot of gene expression reconstruction accuracy from mapping single-cell gene signatures to spatial.

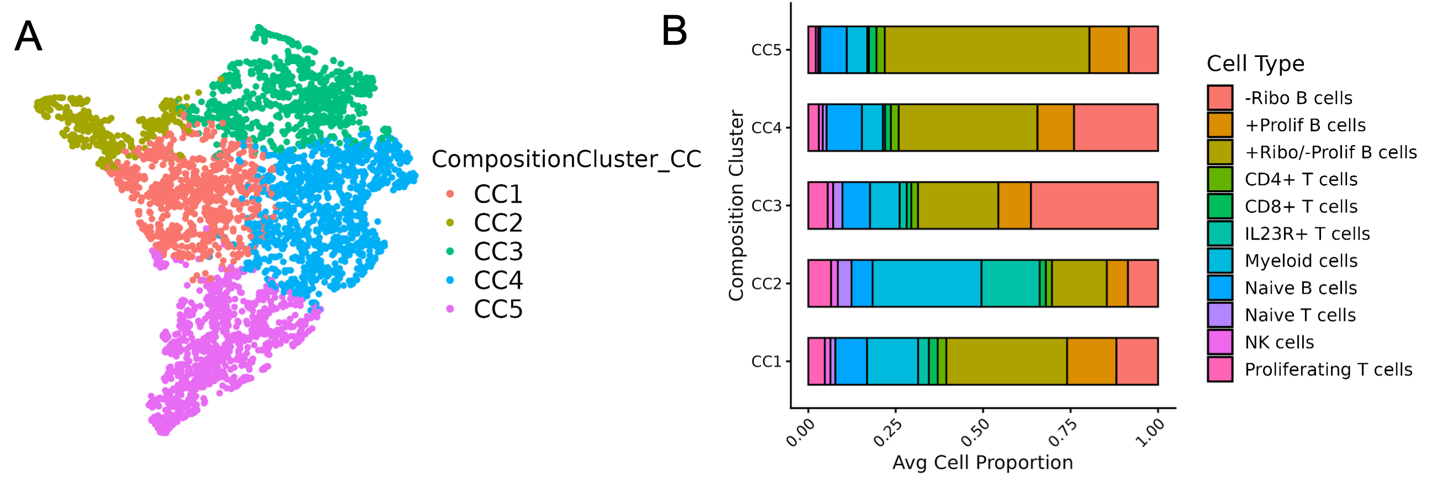

**Supplementary Figure 10. +Ribo/-Prolif B cells and myeloid cells drive cell type stratification of composition clusters. A)** UMAP of composition clustering results at k=5. **B)** Bar graph of composition cluster cell type profiles.
